# Glycine-Rich RNA-Binding Protein 7 interacts with and potentiates effector-induced immunity by Gpa2 and Rx1 based on an intact RNA Recognition Motif

**DOI:** 10.1101/2021.05.18.444452

**Authors:** Octavina C. A. Sukarta, Qi Zheng, Erik J. Slootweg, Mark Mekken, Melanie Mendel, Vera Putker, Hein Overmars, Rikus Pomp, Jan Roosien, Sjef Boeren, Geert Smant, Aska Goverse

**Author notes:** These authors have equal contributions to this manuscript (Octavina C. A. Sukarta, Qi Zheng). To whom correspondence should be addressed: Aska Goverse.

## Abstract

- The activity of intracellular plant Nucleotide-Binding Leucine-Rich Repeat (NB-LRR) immune receptors is fine-tuned by interactions between the receptors and their partners. Identifying NB-LRR interacting proteins is, therefore, crucial to advance our understanding of how these receptors function.
- A Co-Immunoprecipitation/Mass-Spectrometry screening was performed in *Nicotiana benthamiana* to identify host proteins associated with the Gpa2 CC-NB-LRR, which confers resistance against the potato cyst nematode *Globodera pallida*. A combination of biochemical, cellular, and functional assays was used to assess the role of a candidate interactor in defence.
- A *N. benthamiana* homolog of the Glycine-Rich RNA-Binding Protein 7 (*Nb*GRP7) protein was prioritized as a novel Gpa2-interacting protein for further investigations. *Nb*GRP7 also associates *in planta* with the homologous Rx1 receptor, which confers immunity to Potato Virus X. We show that *Nb*GRP7 positively regulates extreme resistance by Rx1 and cell death by Gpa2. Mutating the *Nb*GRP7 RNA recognition motif compromises its role in Rx1-mediated defence. Strikingly, ectopic *Nb*GRP7 expression impacts the steady-state levels of Rx1, which relies on an intact RNA recognition motif.
- Combined, our findings illustrate that *Nb*GRP7 is a novel pro-immune component in effector-triggered immunity by regulating Gpa2/Rx1 functioning at a post-transcriptional level.

## INTRODUCTION

The plant innate immune system is orchestrated by a consortium of cell-autonomous receptor proteins (Bezerra-Neto *et al*., 2020). On the cell surface, Pattern Recognition Receptors (PRRs) detect conserved Pathogen-Associated Molecular Patterns (PAMPs) or damage inflicted on the host (Danger-Associated Molecular Pattern or DAMPs) (Jones & Dangl, 2006; Zipfel, 2014). This triggers basal defence coined as PAMP-triggered immunity (PTI). Pathogens, however, can adapt by evolving virulence-promoting effector molecules to disarm PTI and/or interfere with other host cellular processes (Jones & Dangl, 2006). In this interplay, plants evolved Resistance proteins (R proteins), the majority of which belongs to the family of Nucleotide-Binding Leucine-Rich Repeat (NB-LRR) receptors (Bezerra-Neto *et al*., 2020). Classical NB-LRRs modules have a tri-domain architecture consisting of a central Nucleotide-Binding APAF-1, R-Protein, and CED4 (NB-ARC) region flanked by a N-terminal domains (typically a coiled-coil (CC) or Toll/Interleukin Receptor-like (TIR) domain) and a C-terminal Leucine-Rich Repeat (LRR) domain (van der Biezen, E. A. & Jones, J. D., 1998a, b; Jones *et al*., 2016). NB-LRRs act as a molecular switch that can readily toggle between ADP-bound inactive and ATP-bound active states (Takken *et al*., 2006). The switch function is triggered by recognition of race-specific effector molecules to trigger Effector-Triggered Immunity (ETI). ETI can effectively limit pathogen ingress and is often hallmarked by the visible sign of programmed cell death (Balint-Kurti, 2019). However, the sequence of events leading to immunity remains largely unresolved.

Plant NB-LRRs engage in various interactions with other components in the host proteome, either as preformed complexes or as an active response to a pathogenic intrusion (Sun *et al*., 2020). The common view is that these interactions modulate immunity by regulating defence signalling and/or affecting the stability, localization or activity of the receptor (Sacco *et al*., 2009; Sukarta *et al*., 2016; Sun *et al*., 2020; van Wersch *et al*., 2020). In a vast majority of cases, binding to these co-factors is mediated by domains at the receptor’s N-terminal end (Sun *et al*., 2020). This is also consistent with reports showing that the CC/TIR domains of a few NB-LRR systems can multimerize upon activation, which is thought to increase the surface area available for scaffolding interacting partners (Bentham *et al*., 2018). The nature of proteins known to bind to an NB-LRR varies, ranging from well-established molecular-chaperones (e.g., SGT1 and RAR1) to transcription factors (Bieri *et al*., 2004; de la Fuente van Bentem *et al*., 2005; Leister *et al*., 2005; Tameling & Baulcombe, 2007; Chang *et al*., 2013; Townsend *et al*., 2018). Aside from a few exceptions, however, a limited number of host proteins are known to directly associate with the NB-LRR N-termini (Sun *et al*., 2020). Additionally, how these interactors contribute to NB-LRR immunity is often not fully understood. Uncovering the identity and functions of these interactors will contribute to advancing our understanding of how NB-LRRs mediate defence.

The CC domain of the potato Rx1 immune receptor, which confers resistance to Potato Virus X (PVX), has been shown to act as a scaffold by recruiting various molecular components (Sacco *et al*., 2007; Tameling & Baulcombe, 2007; Townsend *et al*., 2018; Sukarta *et al*., 2020). In the cytoplasm, Rx1 forms a complex with RanGTPase-Activating Protein 2 (RanGAP2) to retain a subpopulation of the receptor in this compartment. This is required by Rx1 to recognize PVX and prompt a complete defence response to the virus (Slootweg *et al*., 2010; Tameling *et al*., 2010). A pool of Rx1 also resides in the nucleus, where it co-opts and modulates the DNA-binding activity of nuclear-associated proteins such as the Golden2-like transcription factor (GLK1) and DNA-Binding Bromodomain Containing Protein (DBCP) (Townsend *et al*., 2018; Sukarta *et al*., 2020). The recruitment of compartment-specific host proteins is thought to grant Rx1 with distinct cellular functions as a molecular sensor and response factor. The CC domain of Gpa2, which mediates defence against the potato cyst nematode *Globodera pallida*, shares considerable homology with the Rx1-CC (95.7% identity at the protein level). Despite bearing significant similarities, however, the Gpa2-CC has only been reported to associate with RanGAP2 (Tameling & Baulcombe, 2007). Whether Gpa2 shares a more extensive pool of interacting components in the nucleus and/or cytoplasm is unknown. Elucidating this will reveal the degree by which homologous NB-LRR receptors diverge in their signalling components. This will in turn, uncover common, critical points for regulating NB-LRR activity.

In the present study, we identified a *Nicotiana benthamiana* homolog of the Glycine Rich RNA Binding Protein 7 (*Nb*GRP7) as a novel interactor of Gpa2. GRP7s are highly conserved plant proteins involved in RNA processing and have previously been implicated in early and late PTI responses (Lee *et al*., 2012; Nicaise *et al*., 2013; Wang *et al*., 2020). However, the function of a GRP7 homolog in ETI has yet to be reported. Here, we present molecular evidence that *Nb*GRP7 is a pro-immunity component in effector-induced immune responses by Gpa2 and its close homolog Rx1. Substituting a conserved arginine residue in the *Nb*GRP7 RNA Recognition Motif (RRM) compromises its potentiating effects on Rx1-mediated resistance, suggesting that RNA-binding may be crucial for the function of *Nb*GRP7 in NB-LRR-mediated immunity. Additionally, we show that *Nb*GRP7 regulates the steady-state levels of Rx1 transcripts and, as a consequence, proteins in the cell. Our results collectively reveal a novel layer of control on the activity of intracellular NB-LRR immune receptors, like Gpa2 and Rx1, at a post-transcriptional level.

## MATERIALS AND METHODS

### Plasmid construction

Full-length *Nb*GRP7 was isolated from *N. benthamiana* cDNA using gene-specific primers listed in **Supporting Information Table S1** as a NcoI-KpnI fragment by High Fidelity PCR (Promega) according to the manufacturer’s protocol. Purified fragments were initially ligated into pGEMT-easy for sequencing and then sub-cloned into the pRAP vector (Schouten *et al*., 1997) containing the N-terminal 4×Myc.GFP tag by 3-way ligation (1:1:1 ratio) following additional BspHI digestion reactions. Positive clones were finally cloned into the pBINPLUS binary vector (van der Vossen *et al*., 2000) as AscI-PacI fragments in A. tumefaciens MOG101. The full-length nucleotide sequence of *Nb*GRP7 was deposited in Genbank with accession MW478352.

For targeted substitution of *Nb*GRP7 R49Q and R49K, nested PCR was performed using primers listed in **Supporting Information Table S1** and Ready-ToGo beads (illustra PuReTaq PCR Beads, *GE Healthcare*). In the first round, primers were used to amplify regions encompassing the mutation in the RNA recognition motif. The resultant fragment was used as template in a second round of PCR with overlapping extensions to obtain the full-length *Nb*GRP7 fragment. The same cloning steps for addition of 4×Myc.GFP tag and into the binary pBINPLUS vector was performed as listed above.

For hairpin silencing, potential silencing regions in *Nb*GRP7 were screened using the Solgenomics VIGS tool (http://solgenomics.net/tools/vigs) against the *N. benthamiana* gene models database v.04.4. Selection of optimal regions included least probability of off-target effects. Target sequences were ordered synthetically (Genescript) in antisense orientation with a spacer in between as specified in **Supporting Information Table S2**. These were subcloned into the destination vector pPT2 (Shin *et al*., 2017) by BamHI/XbaI digestion first in *E. coli* TOP10 and finally, *A*.*tumefaciens* strain MOG101.

For Bi-Fluoresecnce complementation (Bi-Fc), NbGRP7, Rx1-CC and Gpa2-CC were cloned initially into pENTR-D topo vector (Invitrogen). Sequence-verified fragments were then cloned into both pDEST-SCYNE(R)^GW^ or pDEST-SCYCE(R)^GW^ vectors by Gateway LR reaction as described (Gehl *et al*., 2009; Diaz-Granados *et al*., 2020).

### *Agrobacterium tumefaciens* transient assay (ATTA)

ATTA was used as a system for heterologous protein expression in plants as described in (Slootweg *et al*., 2010). Final agrobacterial suspensions were diluted to final OD_600_ values according to each assay. Agroinfiltration was performed on the underside of the leaves of 2-3 weeks old *N. benthamiana* plants using needleless syringes. Plants were grown under standard glasshouse conditions at a constant temperature of 23°C with light and dark cycle of L18:D6. Infiltrated spots were screened for the development of cell death, harvested for protein extraction or examined by microscopy at 1-5 days post infiltration (dpi) depending on the assay and construct.

### Protein extraction and immunodetection

Protein extraction was performed as described in (Slootweg *et al*., 2010). Briefly, 50-100 mg of leaf material was grounded in extraction buffer (10 mM DTT, 150 mM NaCl, 50 mM Tris-HCl, pH 7.5, 1 mM EDTA, 10% glycerol, 2% polyvinylpolpyrrolidone, and 0.5 mg/mL pefabloc SC protease inhibitor [Roche]), and spun down at 16,000 rpm for 5 minutes at 4°C. The supernatant was run through a G25-sephadex column and the eluate was used for subsequent pull-down assays or mixed directly with 4X Nupage LDS sample buffer with 1M DTT (Invitrogen). Proteins extracted were then separated by loading onto 12% Sodium dodecyle sulfate-Polyacrylamide gel electrophoresis (SDS-PAGE) run in 1X MOPS buffer and visualized by Commassie Brilliant Blue staining or wet blotting. Myc-tagged candidate interactors were detected using Goat α-Myc polyclonal antibodies (Abcam) in subsequent western blot analysis as described by (Tian *et al*., 2014). However, hereby immunodetection was achieved using a second polyclonal antibody conjugated with Horse-Radish Peroxidase (Abcam). Conversely, HA and GFP-tagged fusion proteins were detected using a Peroxidase-conjugated α-HA (Roche) or α-GFP (Abcam) antibodies respectively. Finally, peroxidase activity was detected by reacting with the Dura luminescenet and SuperSignal West Femto substrates (1:1 ratio; Thermo Scientific, Pierce) using the G:Box gel documentation system (Syngene).

### *In planta* Co-Immunoprecipitation assays (Co-IP)

N-terminally tagged constructs for expression of p35S:Rx1-4×HA.GFP, p35S:Rx1 S1-4×HA.GFP, p35S:Rx1 S4-4×HA.GFP, p35S:4×HA-Rx1 CC S1, p35S:4×HA-Rx1 CC S4, p35S:4 × HA-Rx1.CC, p35S:4 × HA-Gpa2.CC, p35S:4 × HA.Gpa2 and p35SLS:4×HA.GFP were as described in (Slootweg *et al*., 2010) and (Slootweg *et al*., 2018). *N. benthamiana* leaves infiltrated by the appropriate protein combinations (at OD_600_ of 0.3-0.5) were harvested at 2-3 dpi. For Co-IP, proteins were extracted as described above. Prior to the pull-down, protein samples were pre-cleared by incubation with mouse IgG1 agarose beads. After mixing with α-GFP, α-Myc or α-HA magnetic beads (μMACS) and washing, eluted proteins were run in an SDS-PAGE system (Bis-Tris gel, 12%, Invitrogen) with 1X MOPS buffer and blotted onto PVDF membrane. Immunodetection was then performed as described beforehand using the appropriate antibodies.

### Co-Immunoprecipitation/Mass-Spectrometry analysis

For proteomics analysis, p35S:Gpa2.CC-GFP or p35S:GFP was expressed transiently in *N. benthamiana* between 22-28 hours. Proteins were extracted from leaf samples according as detailed previously and used in cell fractionation as described in (Slootweg *et al*., 2010). Bait proteins were precipitated using μMACS α-GFP beads (Miltenyi) as described above. Peptides were generated by on-beads trypsin digestion of the pull-down samples, which were subsequently sent for MS analysis at the Proteomics Centre at WUR Biochemistry (Wageningen). For identification of proteins, the spectra of each run was matched using a MaxQuant software via a database consisting of translated ESTs and UniProt data referring to *N. benthamiana* and *N. tabacum*.

### Confocal laser scanning microscopy

Cellular localization studies were performed using the Zeiss LSM 510 or the Leica SP8-SMD confocal microscope (for BiFc experiments). Agrobacteria harboring the appropriate constructs were infiltrated on *N. benthamiana* leaves at final OD_600_ values of 0.3-0.5. Leaf epidermal cells were harvested at 2-3 dpi for imaging as described previously in (Slootweg *et al*., 2010). For BiFc measurements, the white laser was used to excite SCFP3A and chlorophyll auto-fluorescence at 440nm and 514nm, respectively. SCFP3A was detected at emission wavelength of 448nm to 495nm. Chlorophyll auto-fluorescence was detected at emission wavelength of 674nm to 695nm. Analysis of fluorescence intensities was performed using the ImageJ application software.

### Chlorophyll assay

Chlorophyll content was measured to indicate degree of cell death as described previously in (Harris *et al*., 2013). Briefly, 3 mm discs of infiltrated *N. benthamiana* leaves were incubated overnight in DMSO at 37°C with constant rotation (250 rpm). Subsequently, absorption measurements of the DMSO solution was read at wavelengths 450 nm and 655 nm using the BioRad Microplate Reader (model 680). Uninfiltrated leaf discs were used as negative controls.

### PVX resistance assay

Viral accumulation was quantified using DAS-ELISA as described in (Slootweg *et al*., 2010). Briefly, plates were coated with polyclonal antibodies (1:1000) raised against the viral CP (Prime Diagnostics). A second polyclonal antibody conjugated with alkaline phosphatase was used for immunodetection (1:1000) at wavelength 405 nm (BioRad Microplate Reader model 680) via the substrate p-nitrophenyl-phosphate. Absorbance Measurements were taken with a reference filter of 655 nm.

### Expression analysis by qRT-PCR

Total RNA was extracted from 50 mg leaf tissues using the Promega Maxwell 16 simpleRNA extraction kit according to the manufacturer’s protocol. First-strand cDNA synthesis was directly performed using the SuperScript III First-Strand Synthesis System (Invitrogen). To analyze expression levels, qRT-PCR was done (BioRad System) in a total reaction mix of 25 µl consisting of: 1 µl forward and reverse primers (5 mM each), 8.5 µl Taq ready mix and 12.5 µl MQ water. qPCR was run using the following program: initial denaturation at 95°C for 15 min followed by 40 cycles of amplification at 95°C for 30s, 60°C for 30s, 72°C for 30s and final elongation at 72°C for 60s with a 90X melting curve at 50°C for 10s. To promote reproducibility, each sample was analyzed *in duplo*. In addition, a standard no template control was included to indicate the presence of contaminating DNA. qPCR data was normalized against the actin housekeeping gene. Finally, relative expression levels were analyzed by the comparative method (2^-ΔΔCt^) using the average threshold values as described in (Schmittgen & Livak, 2008).

### Statistical test

Statistical analyses was performed in R studio Version 1.1.456. Data from assays performed in this study were checked for normality using the Shapiro-Wilk Test. Depending upon the outcome of the normality test, statistical level was determined either by T-test or Wilcoxon-signed rank test with α = 0.05.

## RESULTS

### Identification and isolation of *Nb*GRP7 as a Gpa2-interacting protein

To screen for putative interactors of the Gpa2-CC domain, we adopted a targeted proteomics approach by performing cellular fractionation coupled with Co-IP/MS analysis in *N. benthamiana*. To that end, Gpa2-CC-GFP or GFP (negative control) bait constructs were generated under the control of the Cauliflower Mosaic Virus (Cam35VS) promotor for transient overexpression in *N. benthamiana* by ATTA. As anticipated, MS analysis of the eluted fractions showed an overrepresentation of the Gpa2-CC-GFP bait in both cellular extracts. We also co-purified RanGAP2 exclusively in the cytoplasmic fraction of the pull-down consistent with previous studies (Sacco *et al*., 2007; Tameling & Baulcombe, 2007). This finding supports that the technical approach and stringency used for the data analysis were sound. Interestingly, a protein that has significant peptide hits matching to a GRP7 homolog co-precipitated consistently with the Gpa2-CC nuclear fraction (**Supporting Information** Fig. **S1**). Given the specificity and reproducibility of the interaction observed, we prioritized GRP7 in further studies as described below.

To facilitate functional analysis, we isolated the predicted *N. benthamiana* GRP7 homolog (*Nb*GRP7) based on the *N. benthamiana* draft genome (Solgenomics) and existing *At*GRP7 sequence. The isolated transcript is 501 bp long, encoding a protein of 167 amino acids with an estimated weight of ∼16.9 kDa. Additional sequence alignment showed that *Nb*GRP7 exhibits strong similarity with other plant-derived GRPs, sharing the highest sequence identity (73-75% at the amino acid level) to the *Solanum tuberosum* GRP7 variants (XP_006365106.1 and XP_006365107.1) (**Supporting Information** Fig. **S2a**). The *Nb*GRP7 N-terminus constitutes the most conserved region, wherein the canonical RNA Recognition Motif (RRM) resides (**Supporting Information** Fig. **S2b**). Positioned within this region are also two Ribonucleoprotein motifs (RNP1 and RNP2) and arginine residues required for RNA binding (summarized schematically in Fig. **1a**) (Fu *et al*., 2007; Nicaise *et al*., 2013). The variable and highly disordered glycine-rich region accounts for the remaining C-terminal half of the protein, which is further interspersed with aromatic amino acids.

**Fig. 1.**
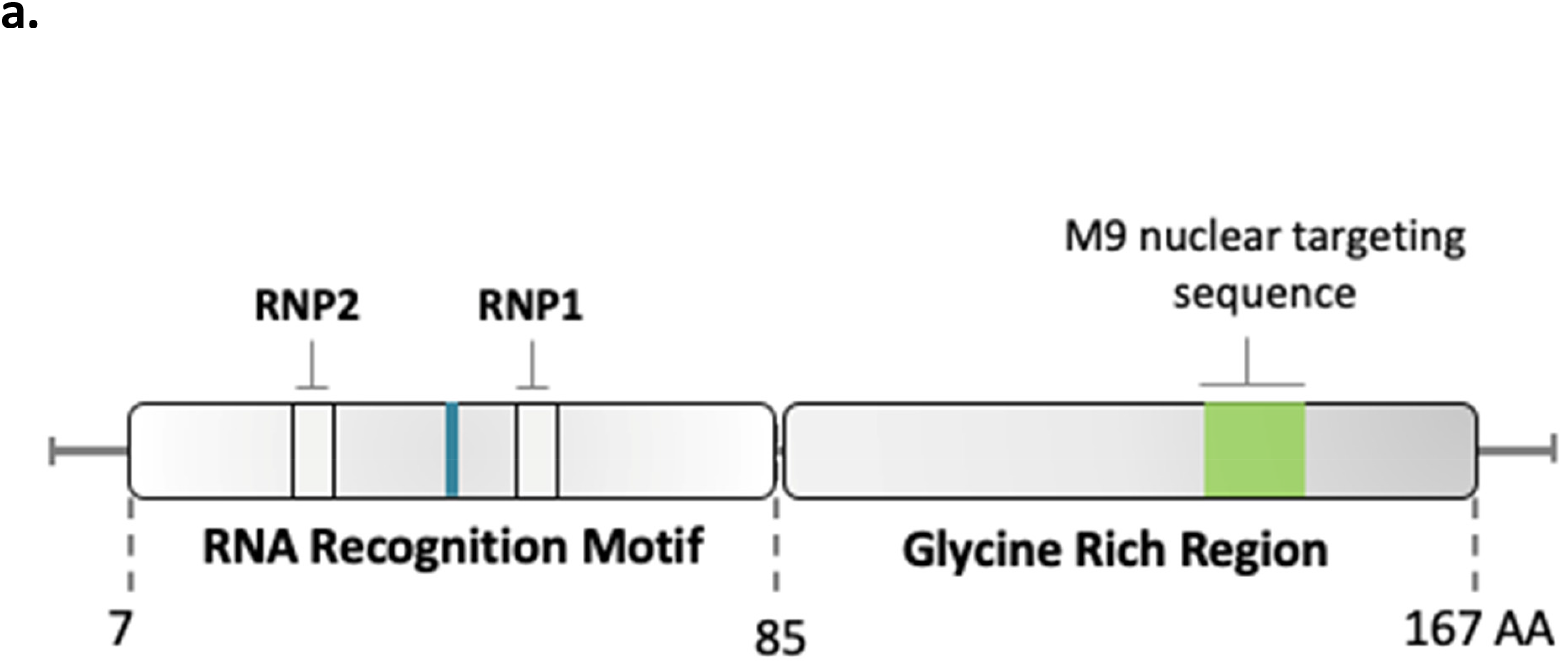
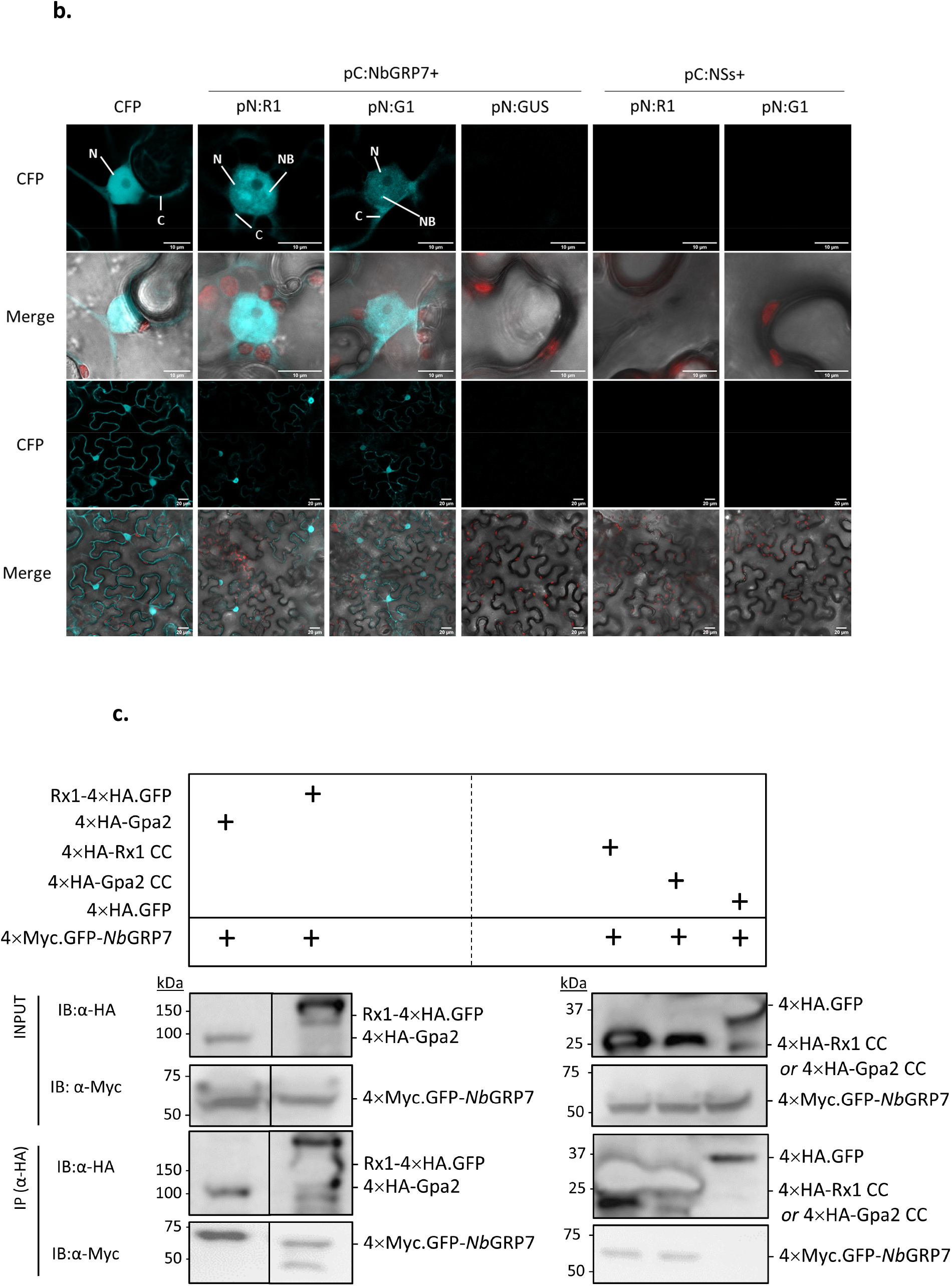
Identification of *Nb*GRP7 as Gpa2 and Rx1-interacting protein. **a)**. Schematic diagram representing the full-length *Nb*GRP7 homolog isolated from *N. benthamiana* cDNA. The conserved Arginine residue required for RNA binding is highlighted in blue. RNP1 = ribonucleoprotein motif 1; RNP2 = ribonucleoprotein motif 2. **b)**. Biomolecular fluorescence complementation (BiFC) of SCFP3A. The N-terminal half of the super cyan fluorescent protein SCFP3A fused Rx1-CC or Gpa2-CC (pN:R1 or pN:G1) and the C-terminal half of SCFP3A fused NbGFP7 (pC:NbGRP7) were co-expressed in *N. benthamiana* leaves. SCFP3A was detected in CFP channel, chlorophyll auto-fluorescence was shown together with CFP signal in the merged channel. Free CFP was used as positive control and co-expression with pC:NSs or pN:GUS were used as negative controls. N, nucleus. C, cytoplasm. NB, nuclear body. Scale bar=10 or 20 μm. Cells were imaged at 2 dpi based on 3 cells. Results are representative of 2 biological repeats. **c)**. Immunoblots from Co-IP of *Nb*GRP7 with full-length Gpa2/Rx1 or their CC domains. For the pull-downs, crude extracts of *N. benthamiana* co-expressing the appropriate protein combinations were incubated with α-HA magnetic beads (μMACS). Gpa2/Rx1 constructs or the negative 4×HA.GFP control were used as baits to co-purify 4×Myc.GFP-*Nb*GRP7.

### *Nb*GRP7 interacts with full-length Gpa2 and Rx1 *in planta* via the CC domain

We next sought to confirm the interaction identified in the Co-IP/MS screening by performing BiFC imaging. Although *Nb*GRP7 was originally found to associate with the Gpa2-CC domain, we expanded our assay to test the interaction of *Nb*GRP7 with Rx1 given its close homology, particularly in the CC which only differs in six amino acid residues. To that end, we created both Gpa2-CC and Rx1-CC constructs fused to the N-terminal half of the super cyan fluorescent protein SCFP3A (pN:G1 and pN:R1) for transient co-expression with *Nb*GFP7, which was fused to the C-terminal half of SCFP3A (pC:*Nb*GRP7). The reverse combinations (pC:R1, pC:G1, and pN:*Nb*GRP7) were also generated for comparison. Combinations co-expressing pN:R1 or pN:G1 with the viral protein NSs (pC:NSs) were used as negative controls. Likewise, the combination of *Nb*GRP7 with β-Glucuronidase (pN:GUS) was used as an additional negative control. Confocal imaging at 2 dpi shows that a CFP signal accumulated in the nucleus and to a lesser extent, in the cytoplasm when either pN:G1 or pN:R1 was co-expressed with pC:*Nb*GRP7 (Fig. **1b**). Remarkably, detailed imaging of the nuclei showed the nucleoplasm to have a non-homogenous distribution with CFP signals accumulating in subnuclear bodies similar to those described in earlier studies of *At*GRP7 (Kim *et al*., 2008). These cellular structures are typically associated with RNA processing, which coincides with the expected function of a GRP7 homolog (Spector & Lamond, 2011). Conversely, a CFP signal was absent upon co-expression of the negative control combinations (pN:R1/pN:G1 with pC:NSs and pC:*Nb*GRP7 with pN:GUS) (Fig. **1b**). All aforementioned constructs were expressed stably and similar results were obtained with the reverse combinations (**Supporting Information** Fig. **S3b**). Combined, our findings confirm that *Nb*GRP7 can form a complex with the CC domains of Gpa2 and Rx1 *in planta*, predominantly in the nucleus.

We next examined whether *Nb*GRP7 can bind to full-length Gpa2 and Rx1 *in planta* by Co-IP. Thus, a 4×Myc.GFP-*Nb*GRP7 construct was co-expressed transiently in combination with (HA)-tagged version of the full-length receptors (4×HA-Gpa2 and Rx1-4×HA.GFP). Infiltrated leaf materials were harvested at 2 dpi and the extracted proteins were subjected to a Co-IP using the α-HA magnetic beads system (μMACS). Immunoblotting of the eluates shows that 4×Myc.GFP-*Nb*GRP7 co-precipitates with both 4 × HA-Gpa2 and Rx1-4 × HA.GFP (Fig. **1c**). Consistent with the Co-IP/MS screening, we also observed 4×Myc.GFP-*Nb*GRP7 to specifically co-purify with the CC-domain of Gpa2 (4×HA-Gpa2-CC). Taken together, our data demonstrate that *Nb*GRP7 protein interacts with full-length Gpa2 and Rx1 immune receptors *in planta*.

Given that *Nb*GRP7 associates with full-length Gpa2/Rx1 and their CC domains, we questioned whether additional receptor domain(s) can contribute to this complex formation. Thus, Co-IP studies were performed using 4×Myc.GFP-*Nb*GRP7 as bait with HA or GFP-tagged fusions of the Gpa2/Rx1 CC, NB-ARC and LRR domains. Interestingly, 4×Myc.GFP*-Nb*GRP7 exclusively co-purifies with 4×HA-Rx1-CC but not the other receptor domains nor the 4×HA.GFP control (**Supporting Information** Fig. **S4a**). This shows that the CC is required and sufficient for the interaction with *Nb*GRP7. To further localize the structural determinants in the CC required for *Nb*GRP7 binding, we used available S1 and S4 surface mutants of the Rx1-CC domain as described in (Slootweg *et al*., 2018). The S4 mutations disrupt the hydrophobic patch essential for RanGAP2-binding in helix 4 of the Rx1-CC, while the S1 mutations in helix 1 reduce intramolecular binding to the NB-LRR. We showed that 4×Myc.GFP-*Nb*GRP7 co-precipitated with the S1 and S4 mutant variants (CC and full-length) similar to the wildtype control (**Supporting Information** Fig. **4b1 and 4b2**). While the immunoblot shows that Rx1 S1-4 × HA.GFP pulled down at a greater extent compared to the wild-type and S4 derivates (**Supporting Information** Fig. **4b2**), this was not consistent between experimental repeats. These findings suggest that S1 and S4 surface regions of the CC are most likely not involved in complex formation with *Nb*GRP7. Thus, *Nb*GRP7 interacts with Rx1-CC at a surface region distinct from those required for intramolecular interactions and RanGAP2 binding.

### *Nb*GRP7 is a positive regulator of effector-dependent defenses by Gpa2 and Rx1

To ascertain the biological relevance of the interaction observed for *Nb*GRP7 and Rx1/Gpa2, we performed a cell death assay in *N. benthamiana* leaves. Agrobacteria harbouring 4 × Myc.GFP-*Nb*GRP7 were co-infiltrated with Rx1/Gpa2 and their matching effectors, namely the coat protein of PVX strain UK3 (PVX-CP UK3) and GpRBP-1 variant D383-1, respectively. Infiltrated spots were monitored for the progression of cell death within 3-5 dpi by measuring chlorophyll loss. Interestingly, transiently overexpressing 4×Myc.GFP-*Nb*GRP7 potentiates GpRBP-1-induced cell death by Gpa2 (under the control of its endogenous promotor) at 5 dpi as indicated by a greater chlorophyll loss compared to the GFP control (Fig. **2a**). To determine whether the pro-immunity functions of *Nb*GRP7 was effector-dependent, we also included an autoactive p35S:Gpa2 D460V construct. Interestingly, 4 × Myc.GFP-*Nb*GRP7 overexpression does not influence autoactivity by p35S:Gpa2 D460V. These results show that *Nb*GRP7 specifically contributes to effector induced cell death.

**Fig. 2.**
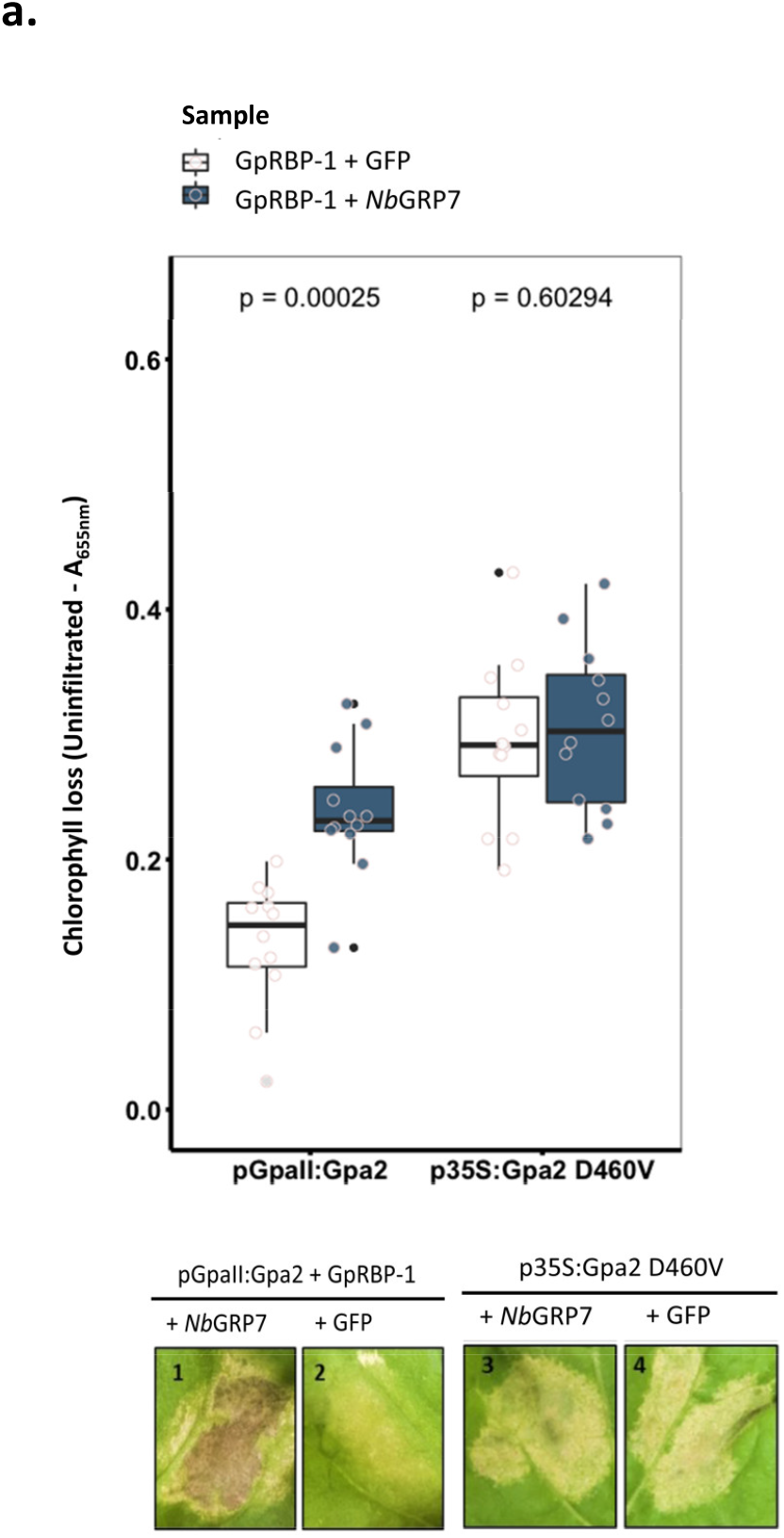
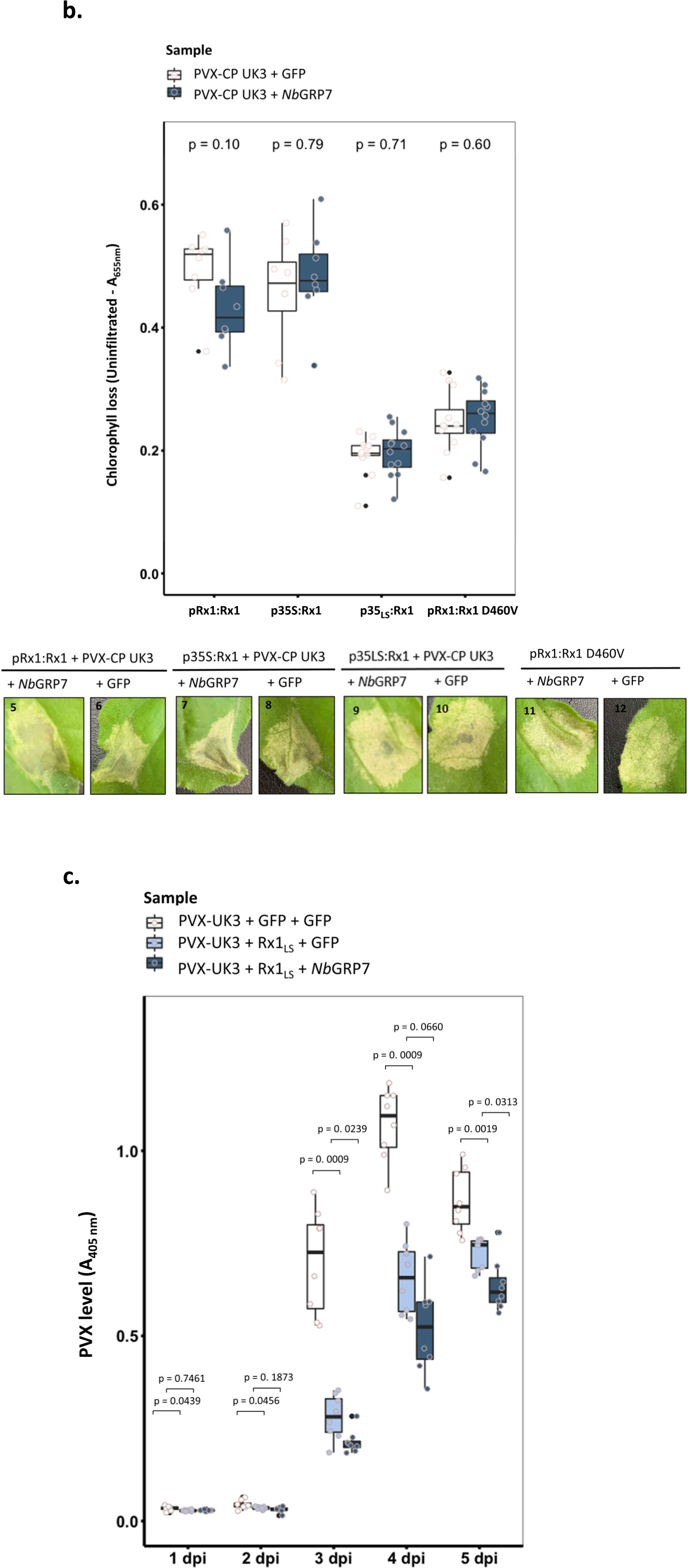
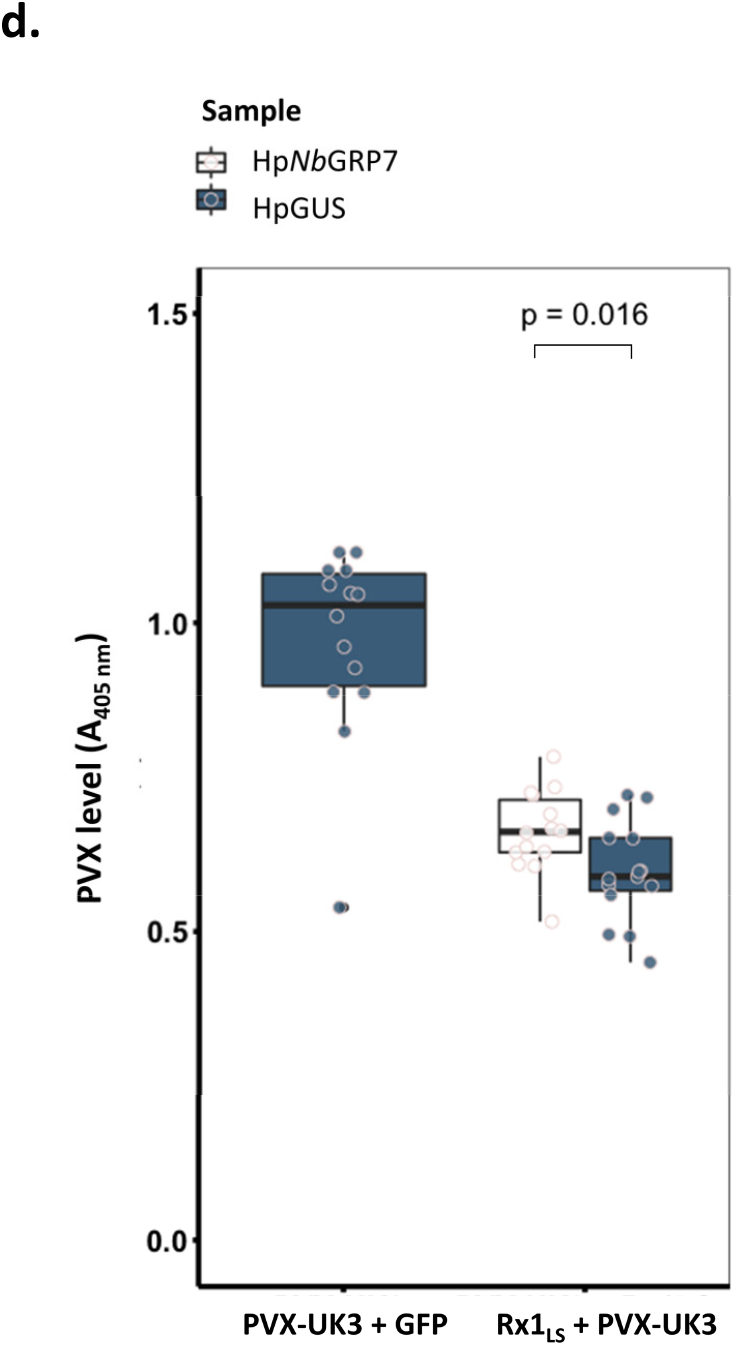
*Nb*GRP7 potentiates defenses by Gpa2 (a) and Rx1 (b,c,d). Boxplots representing chlorophyll loss of elicitor-induced or independent cell death upon overexpression of *Nb*GRP7 at 3-5 dpi (**a, b**). Bars represent the interquartile range while the cross indicates the median. The whiskers mark the minimal and maximal data points. Significance was calculated using Wilcoxon-Signed Rank test with α = 0.05 from n ≥ 12 samples. Data shown is representative of at least three independent repeats. For the cell death assay, constructs of Rx1 under the control of either a 35S, endogenous or leaky scan promotor was used whereas Gpa2 was cloned under the control of its endogenous promotor (pGpaII). Representative photographs of infiltrated leaf zones are provided in the next row. Boxplots of absorbance at 405 nm, indicating levels of PVX-UK3 upon transient overexpression **(c)** or silencing **(d)** of *Nb*GRP7 in the context of Rx1-mediated responses. Data shown is representative of at least three independent repeats with similar results. Significance was calculated using Wilcoxon-Signed Rank test with α = 0.05 from n ≥ 8 samples.

For Rx1, cell death is typically a quick response in *N. benthamiana*. We, therefore, compared cell death induced by Rx1 constructs cloned under the endogenous (pRx1), CaMV35S (p35S), or leaky scan promotor (p35LS as described in (Slootweg *et al*., 2010)). Contrary to Gpa2, transient overexpression of 4 × Myc.GFP-*Nb*GRP7 had negligible effects on Rx1-mediated cell death at 3 dpi (Fig. **2b**). No significant differences in chlorophyll loss relative to the control could be observed reproducibly when overexpressing 4×Myc.GFP-*Nb*GRP7, PVX-CP UK3 and Rx1, irrespective the immune receptor construct used. Likewise, *Nb*GRP7 overexpression did not influence the autoactivity of an pRx1:Rx1D460V construct. Contrary to Gpa2, *Nb*GRP7 does not contribute to Rx1-mediated cell death responses in this study under the conditions used for testing.

Notably, cell death is viewed as a secondary latent response for Rx1 that is reserved by the host when immunity proves insufficient, e.g., when there is an over-abundance of the viral coat protein such as during heterologous expression assays (Bendahmane *et al*., 1999). Instead, PVX infection typically induces an extreme resistance response, which can effectively restrict viral spread without the need to elicit cell death (Bendahmane *et al*., 1995; Bhattacharjee *et al*., 2009). We, therefore, investigated the impact of *Nb*GRP7 overexpression on extreme resistance by Rx1. *N. benthamiana* leaves were infiltrated with Agrobacteria harbouring an amplicon of the avirulent PVX-UK3 strain and p35LS:Rx1. Viral levels were quantified by DAS-ELISA within 1-5 dpi. Our data demonstrate that *Nb*GRP7 enhances Rx1-mediated extreme resistance against PVX-UK3 between 3-5 dpi, as shown by a significantly greater reduction in viral levels compared to the control (Fig. **2c)**. Collectively, these findings indicate that *Nb*GRP7 positively regulates extreme resistance by Rx1. We further show that overexpressing *Nb*GRP7 reduces PVX-UK3 accumulation in the absence of p35LS:Rx1 (**Supporting Information** Fig. **S7**), consistent with existing studies implicating the role of *At*GRP7 in basal defence (Lee *et al*., 2012). These results combined illustrate a role for *Nb*GRP7 in both Rx1-dependent and -independent defences against PVX.

To complement our overexpression studies, TRV-VIGS silencing of *Nb*GRP7 was performed. However, TRV-VIGS silenced plants showed severe developmental phenotypes at 3 weeks post-silencing (data not shown), most likely due to pleiotropic effects of *Nb*GRP7 silencing on accumulation of TRV. We, therefore, abandoned this approach and alternatively, performed local transient hairpin silencing of *Nb*GRP7 (Shin *et al*., 2017). Hairpin constructs (denoted as hp*Nb*GRP7) were designed to knock-down transcript levels of endogenous *Nb*GRP7 specifically upon leaf infiltration (**Supporting Information** Fig. **S5**). Similar p35LS:Rx1 and PVX-UK3 combinations as described above were co-infiltrated with hp*Nb*GRP7, and virus levels were quantified at 3 dpi. The results demonstrate that transient silencing of *Nb*GRP7 leads to significantly higher accumulation of PVX-UK3, indicating that Rx1-dependent resistance was hampered (Fig. **2d)**. These findings complement our overexpression analysis and collectively, support the role of *Nb*GRP7 in extreme resistance by Rx1. Taken together, our data demonstrate that *Nb*GRP7 acts a pro-immune component in Gpa2 and Rx1-meditated effector-dependent defences.

### The function of *Nb*GRP7 in Rx1-mediated extreme resistance depends on an intact RNA Recognition Motif

Having established a role of *Nb*GRP7 in Gpa2 and Rx1 immunity, we questioned whether its capacity to bind RNA may underly the observed phenotypes. Thus, we mutated a conserved arginine residue at position 49 of the *Nb*GRP7 RNA recognition motif to generate mutant variants (*Nb*GRP7-R49K or -R49Q) impaired in their RNA binding as described in (Nicaise *et al*., 2013). Immunoblotting indicates that 4×Myc.GFP-*Nb*GRP7 R49K and 4×Myc.GFP-*Nb*GRP7 R49Q are expressed as stably as wild type 4×Myc.GFP*-Nb*GRP7 (**Supporting Information** Fig. **S6a**). We also compared the subcellular distribution patterns to wild type *Nb*GRP7 using confocal microscopy. Interestingly, the mutant variants showed different cellular distribution patterns as the subnuclear bodies characteristic of *Nb*GRP7 were considerably less prominent (**Supporting Information** Fig. **S6b**).

We then assessed the effects of overexpressing 4 × Myc.GFP-*Nb*GRP7 R49K and 4×Myc.GFP-*Nb*GRP7 R49Q on the Rx1-mediated extreme resistance response as described above. Quantification of virus levels by DAS-ELISA showed that both mutants still potentiate PVX-UK3-induced extreme resistance at 3 dpi, although significantly less than wild-type *Nb*GRP7 (Fig. **3a**). To corroborate these findings, qPCR analysis was performed, which indicates that levels of viral transcripts increased in tissues where p35LS:Rx1 was co-expressed with the mutants compared to wild-type *Nb*GRP7 (Fig. **3b, c**). These results combined suggest that the function of *Nb*GRP7 in Rx1-dependent defences rely on an intact RNA-binding domain.

**Fig. 3.**
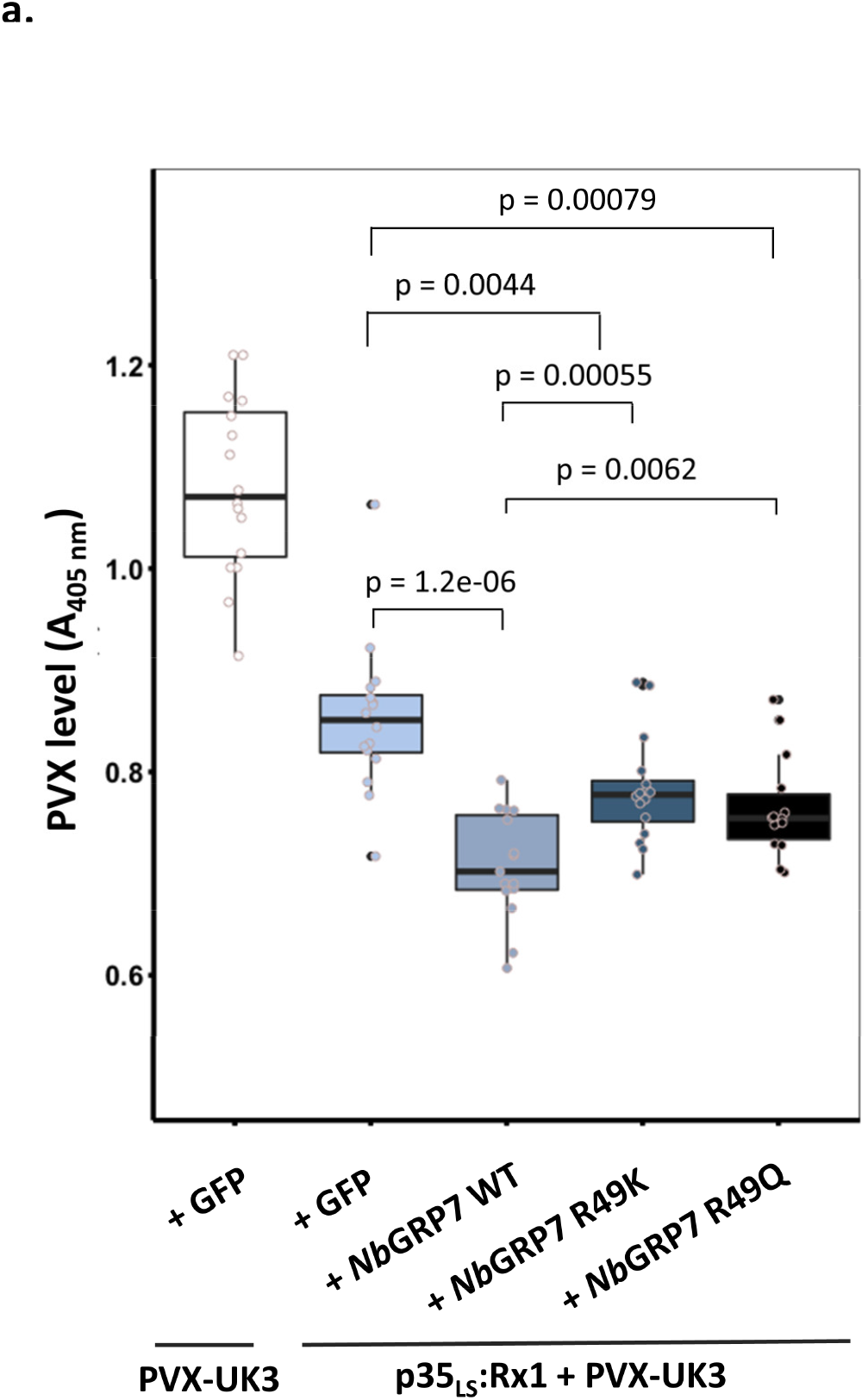
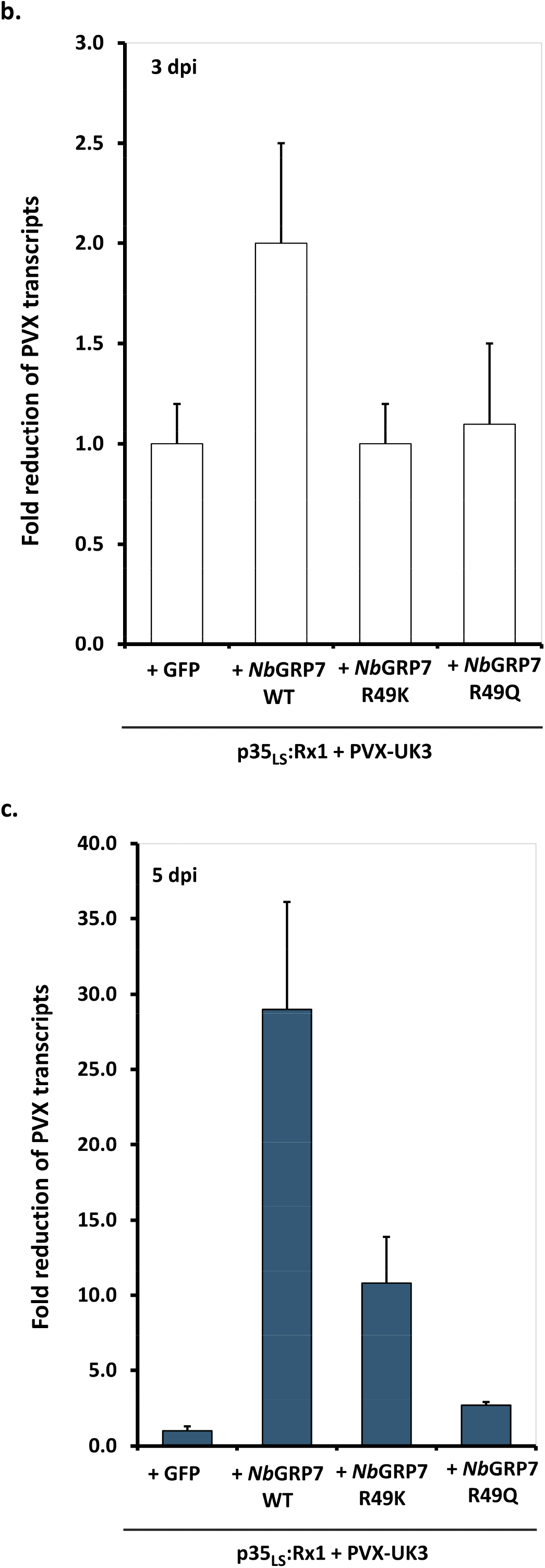
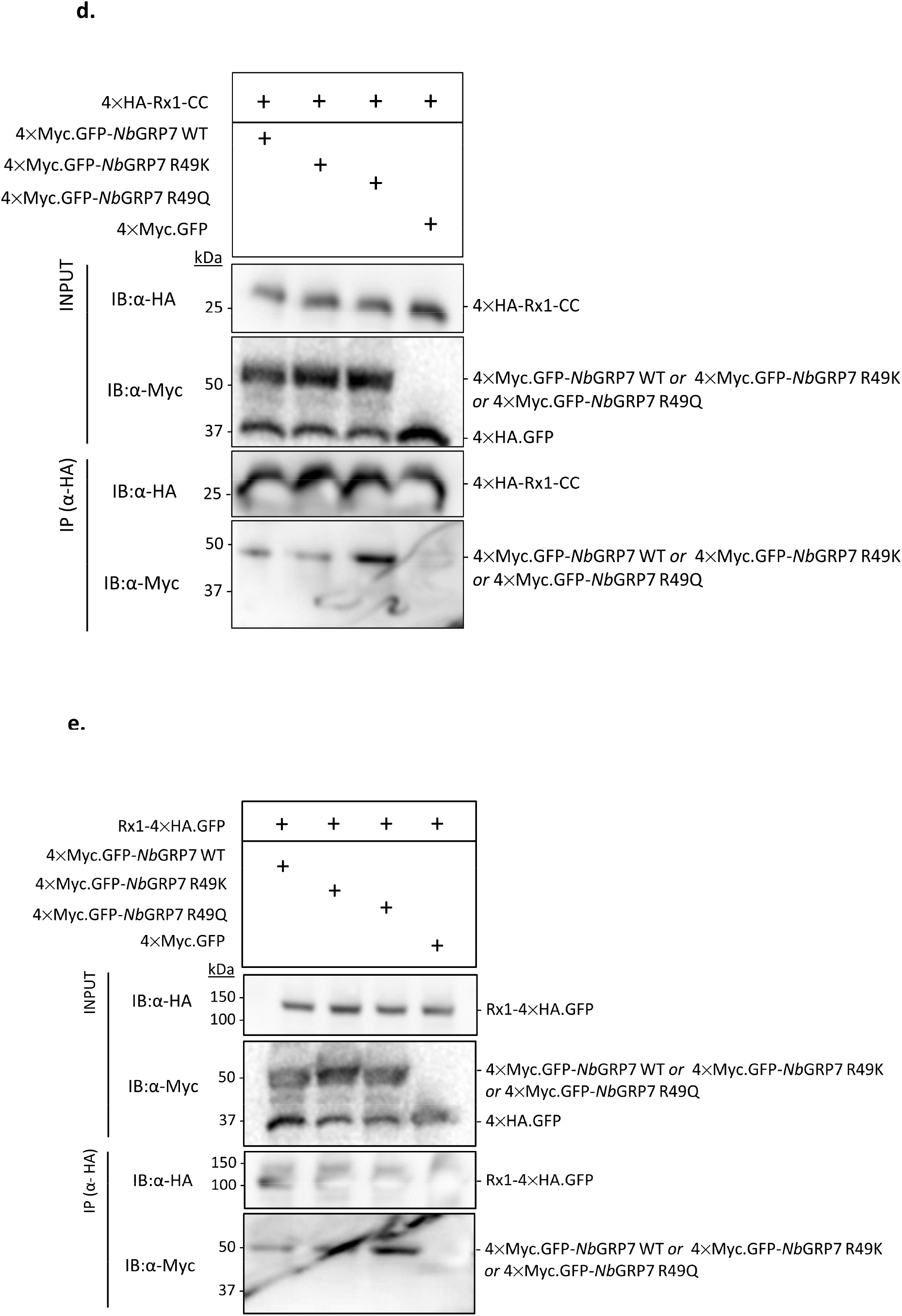
The RNA-binding activity of *Nb*GRP7 contributes to Rx1-mediated extreme resistance against PVX-UK3 in *N*. **a)**. Boxplots of a DAS-ELISA assay from transient overexpression of p35LS:Rx1-GFP and PVX-UK3 in combination with 4×Myc.GFP-*Nb*GRP7 WT, *Nb*GRP7 R49K or *Nb*GRP7 R49Q. Bars represent the interquartile range, and the cross indicates the median. The whiskers mark the minimal and maximal data points. Statistical significance was calculated using Wilcoxon-Signed Rank test with α = 0.05 from n ≥ 12 samples. **b, c)**. Bar graphs of qRT-PCR analysis of viral transcript levels as determined using primers specific for the PVX coat protein. RNA from infected *N. benthamiana* leaves harvested at 3 dpi (**b**) and 5 dpi (**c**) were used for the analysis. Data shown is representative of two different experiments, with each sample consisting of a pool of at least 5 different plants. To obtain the relative fold change, samples were first normalized to the actin reference gene and then compared to the combination of Rx1LS + PVX-UK3 + GFP. Error bars represent standard error. **d, e)**. Co-IP of HA-tagged Rx1-CC domain or the full-length immune receptor in combination with WT or mutated variants of 4×Myc.GFP-*Nb*GRP7. α-HA beads were used to pull-down the receptor fragments. The success of the Co-IP is detected in the α-Myc immunoblot. “+” indicates the presence of a particular construct in the infiltration combination.

To confirm whether the *Nb*GRP7 mutant variants still interact with Rx1, a Co-IP was performed using similar experimental set-ups as described beforehand. Immunoblotting shows that 4×Myc.GFP-*Nb*GRP7 R49K co-immunoprecipitated at comparable levels with 4×HA-Rx1-CC and 4×HA-Rx1 as the wild-type *Nb*GRP7 (Fig. **3d, e**). Thus, we concluded that the reduced pro-immune activity of *Nb*GRP7 R49K is not due to a lack of complex formation with the Rx1-CC, but most likely from the loss of its RNA-binding capacity. Notably, however, the *Nb*GRP7 R49Q variant pulled-down consistently to a greater extent than wild-type *Nb*GRP7. Thus, substituting the conserved arginine residue in *Nb*GRP7 to an amino acid with markedly different properties enhanced its physical interaction with Rx1. Coupled with our functional data, this suggests that the contribution of *Nb*GRP7 in Rx1 defence may also rely on its interaction with the immune receptor.

### *Nb*GRP7 maintains the steady-state levels of Rx1 *in planta*

Our data show that *Nb*GRP7 strengthens Rx1-mediated extreme resistance is dependent on an intact RNA recognition motif. This suggests that the RNA chaperone activity of *Nb*GRP7 may underlie its function in Rx1-mediated defence, for example by the stabilisation of Rx1 transcripts as described for *At*GRP7 and FLS2 (Nicaise *et al*., 2013). To explore this model, we investigated whether overexpressing and silencing of *Nb*GRP7 affects mRNA levels of Rx1 in the cell by performing qPCR analysis. Our findings show that the relative abundance of Rx1 transcripts increased upon *Nb*GRP7 overexpression in the absence of PVX by c.a. 2 to 4-fold when compared to the GFP control at 3 and 5 dpi (Fig. **4a**). Conversely, silencing *Nb*GRP7 decreased Rx1 transcript levels. We reproduced these assays under activating conditions of Rx1 by PVX-UK3. Similar changes in Rx1 transcript profiles were observed during immune activation by PVX-UK3 (Fig. **4b**). These findings combined show that *Nb*GRP7 can modulate the steady-state transcript levels of Rx1 in the cell.

**Fig. 4.**
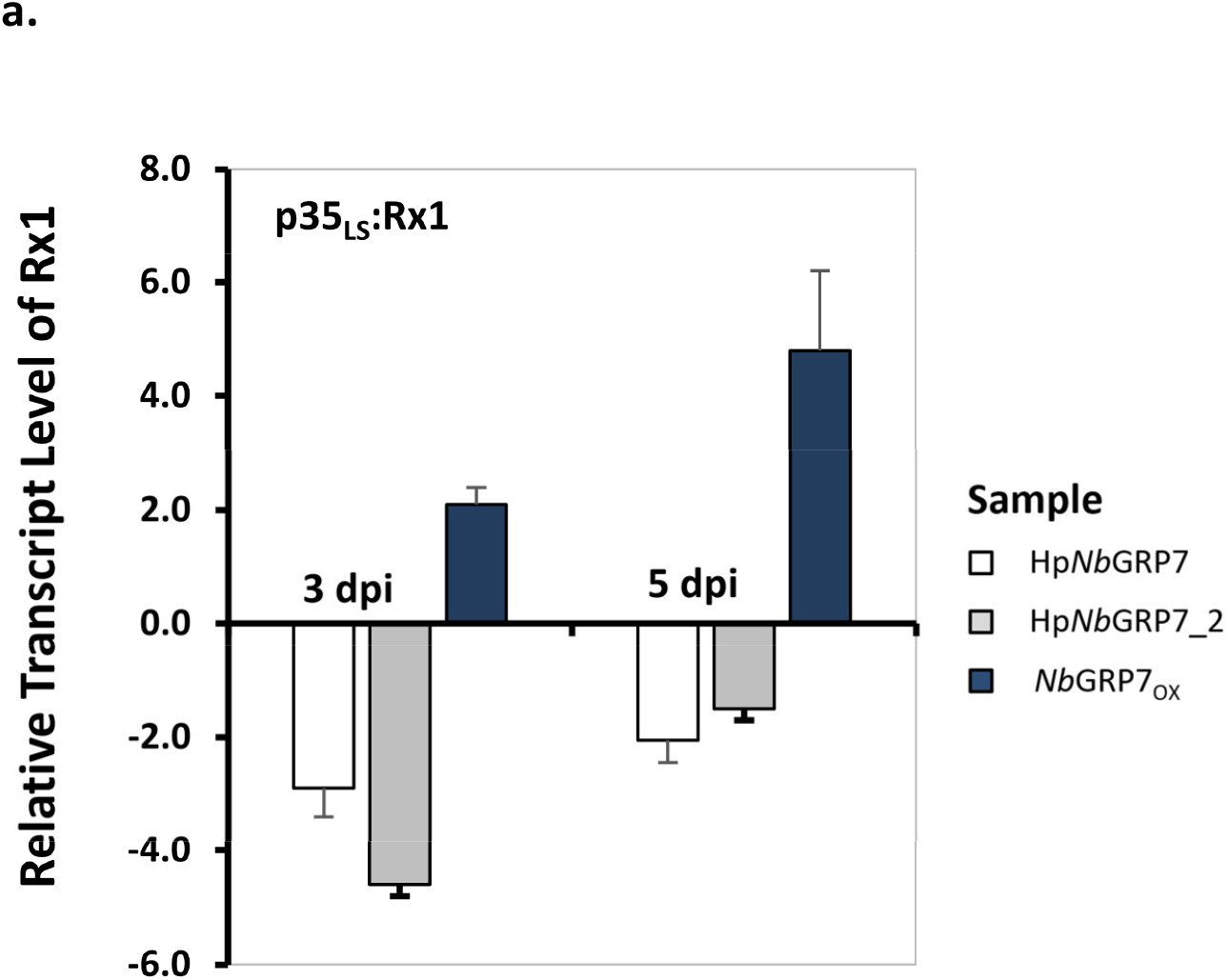
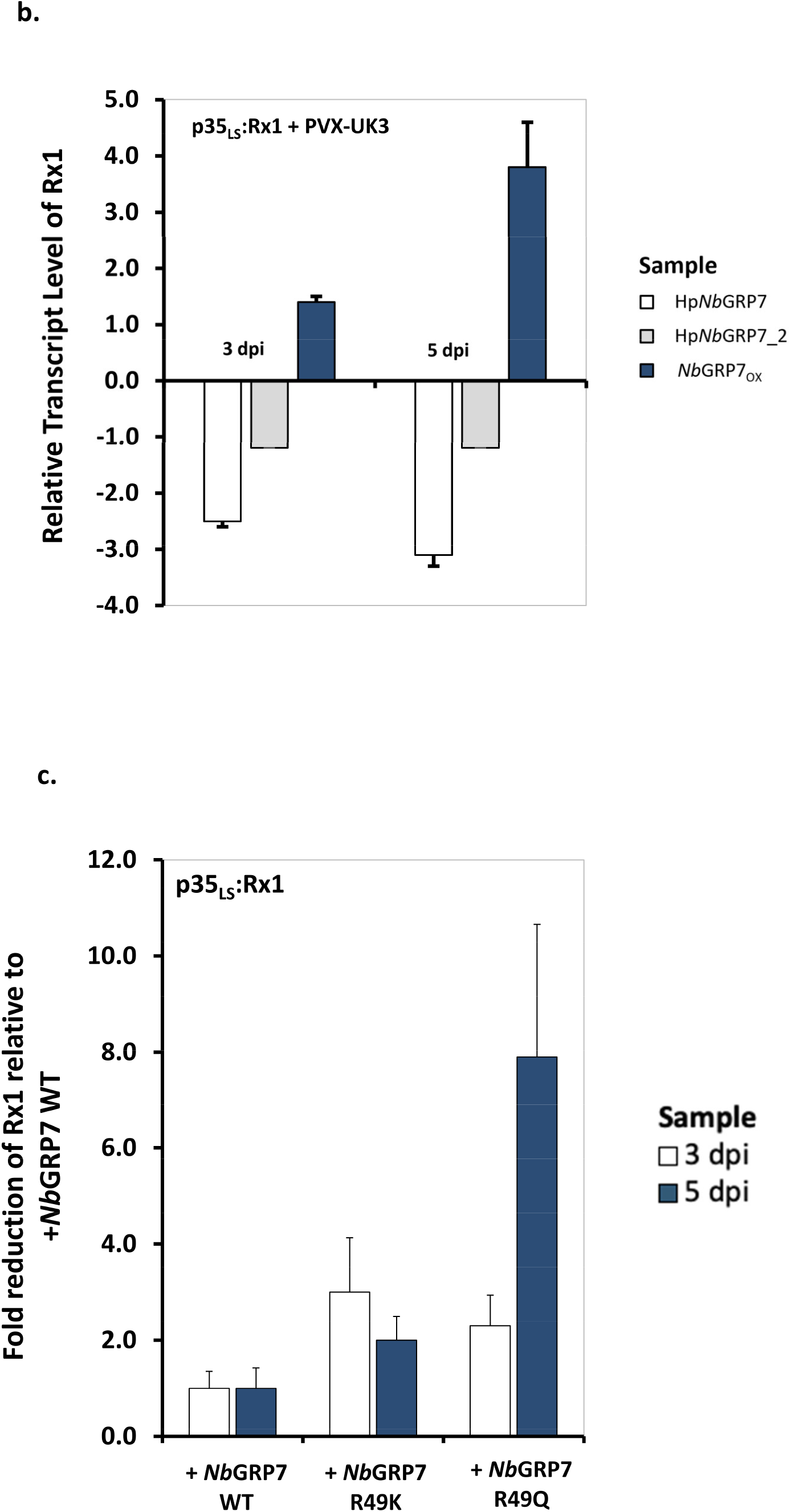
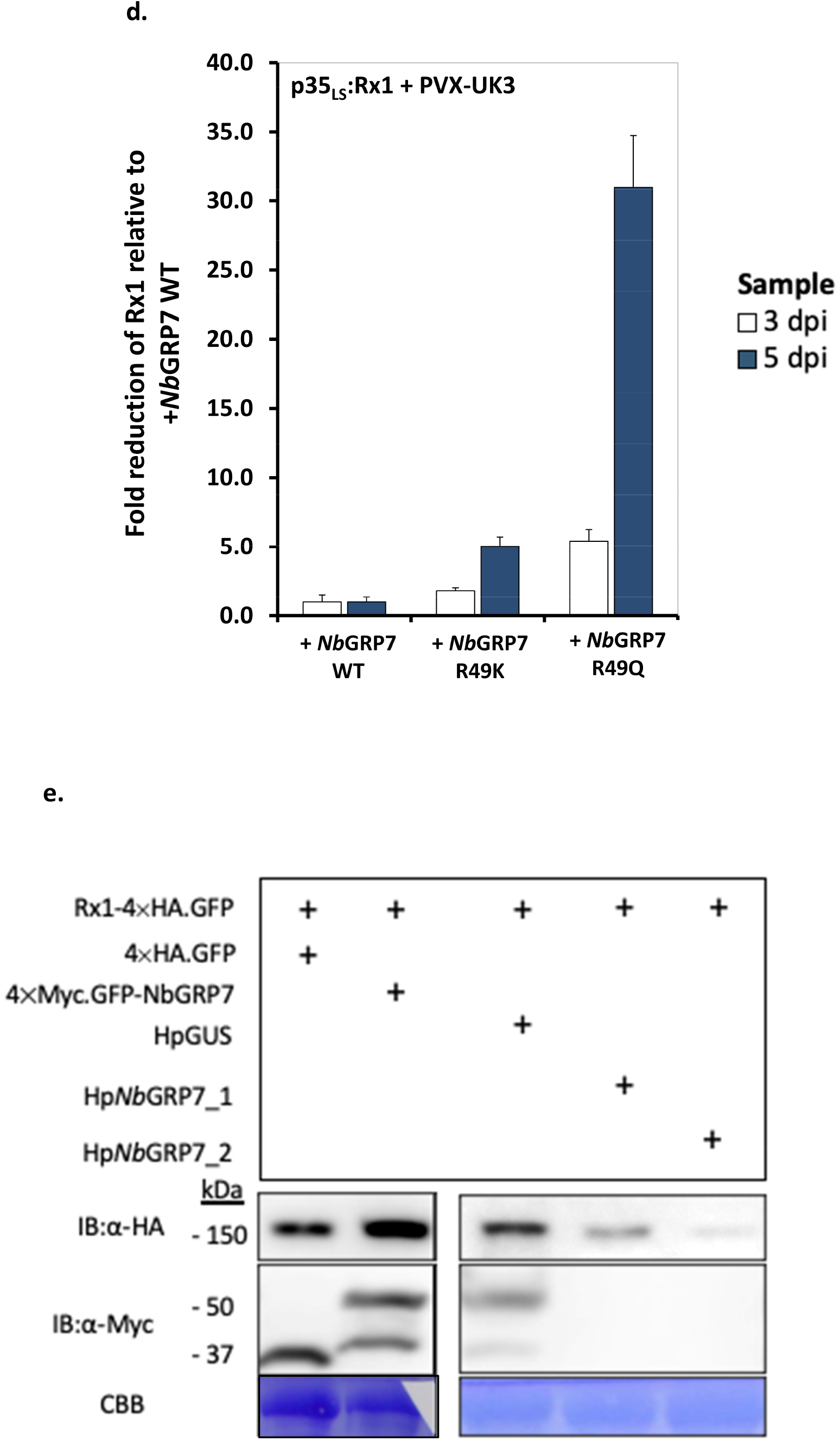
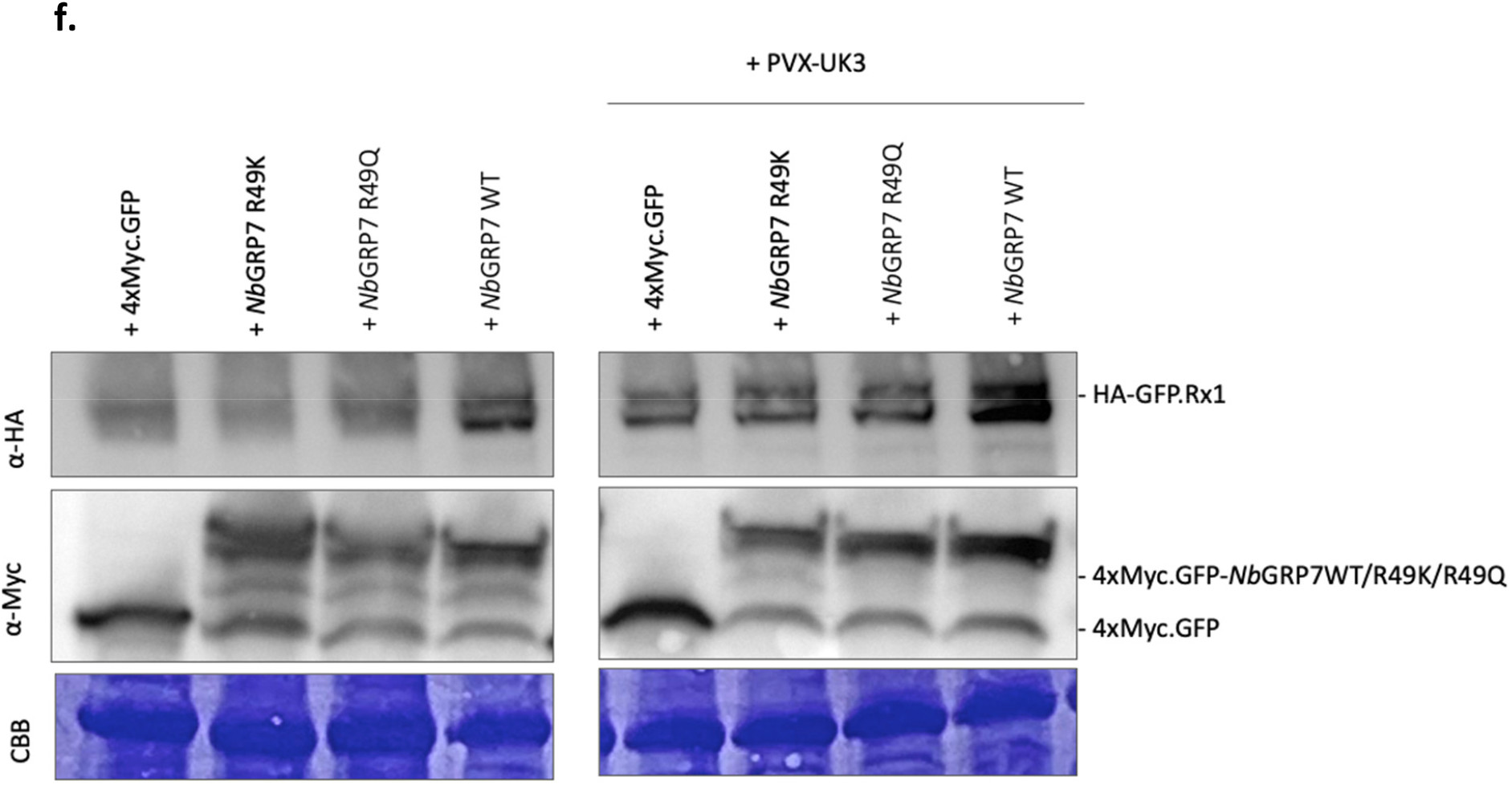
*Nb*GRP7 regulates Rx1 transcript abundance pre- and post-activation by PVX-UK3. **a, b)**. qRT-PCR showing expression profile of Rx1 transcript co-expressed with a construct whereby *Nb*GRP7 is either overexpressed or silenced in *N. benthamiana* upon activation by PVX-UK3 or in the absence of the pathogen. Leaf samples were harvested at either 3 or 5 dpi. For each combination shown, data was obtained from a pool of at least 5 different plants. Rx1 transcript levels were normalized to the actin reference gene and the fold change was calculated relative to HpGUS (for hairpin silencing experiments) or GFP-GUS (for *Nb*GRP7 overexpression experiments). Error bars represent the standard error. **c, d)**. qRT-PCR quantifying the relative transcript abundance of Rx1 pre and post-activation in the presence of wild-type or mutant *Nb*GRP7 constructs at 3 or 5 dpi. Fold change was derived following normalization to the actin reference gene and compared to the combination containing wild-type *Nb*GRP7. Error bars represent the standard error. **e)**. Ectopic expression of *Nb*GRP7 affects the protein abundance of Rx1. Immunblot of protein extracts from *N. benthamiana* leaves co-expressing full-length Rx1 in combination with 4×Myc.GFP, 4×Myc.GFP-*Nb*GRP7 or the hairpin silencing constructs. Leaf samples were harvested at 3 dpi. Data shown is from a single representative experiment. CBB-stained membrane of the RUBisCO protein served as loading control. **f)**. Immunoblot demonstrating the protein stability of full-length HA.GFP-Rx1 in combination with the overexpression of 4×Myc.GFP, 4×Myc.GFP-*Nb*GRP7, 4×Myc.GFP-*Nb*GRP7 R49Q or 4 × Myc.GFP-*Nb*GRP7 R49K. Data shown is from a single representative experiment. CBB-stained membrane of the RUBisCO protein served as loading control.

If *Nb*GRP7 stabilises Rx1 transcripts, we anticipated that this would result in a concomitant increase in Rx1 protein levels. Indeed, immunoblotting assays shows that overexpression of 4 × Myc.GFP-*Nb*GRP7 led to higher protein accumulation of p35LS:GFP-Rx1 while *Nb*GRP7 silencing reduced this amount (Fig. **4e, f**). Moreover, we could demonstrate that this increase in Rx1 transcript and protein abundance depends on the RNA-binding capacity of GRP7. Upon overexpression of the *Nb*GRP7 RNA-binding mutants *Nb*GRP7 R49K and *Nb*GRP7 R49Q, reduced transcript and protein levels of Rx1 were observed when compared to wild-type *Nb*GRP7 both in the presence and absence of PVX-UK3 (Fig. **4c, d**). Collectively, our findings indicate that *Nb*GRP7 stabilizes the steady-state level of Rx1, which could explain its pro-immune activity in Rx1-mediated plant defence as described.

## DISCUSSION

The activity of plant NB-LRRs is regulated by their interaction with components in the plant proteome. However, the identities and functions of NB-LRR-associated proteins are largely unknown. In this study, we describe the identification of *Nb*GRP7 as a novel interactor of the intracellular NB-LRR immune receptors Gpa2 and Rx1 based on a Co-IP/MS screening in *N. benthamiana*. Transient overexpression and silencing experiments demonstrate that *Nb*GRP7 positively contributes to GpRBP-1-dependent cell death by Gpa2 and extreme resistance by Rx1. Interestingly, ectopically expressing *Nb*GRP7 also influenced Rx1 transcript and protein abundance. Both the pro-immune activity and transcript regulation of Rx1 by *Nb*GRP7 rely on an intact RNA-binding domain. Taken together, we infer that *Nb*GRP7 acts as a co-factor regulating the stability of its NB-LRR receptors. We postulate that this occurs at a post-transcriptional level, which is an underexplored mechanism for fine-tuning the functioning of plant NB-LRRs like Gpa2/Rx1. To our knowledge, our research constitutes the first report of the role of a GRP7 homolog in ETI.

By contrast, a role for GRP7 has been explored extensively in the context of basal immunity. For instance, the RNA-binding function of *A. thaliana* GRP7 (AtGRP7) is targeted by the *Pseudomonas syringae* effector HopU1 for ADP-ribosylation to promote virulence of the bacteria (Fu et al., 2007). A more recent study implicates that the phosphorylation of *At*GRP7 induces a dynamic and global alternative splicing response in the Arabidopsis transcriptome upon activation of the FERONIA receptor (Wang *et al*., 2020). Combined with our data, this indicates that PTI and ETI recruit the same pro-immune components present in plant cells to activate defence. This supports the idea that PTI and ETI involve (partial) overlapping pathways in plant immunity. Interestingly, we demonstrate in this study that *Nb*GRP7 can enhance basal resistance against PVX, consistent with the role of *At*GRP7 in FLS2- and Feronia-mediated defences (**Supporting Information** Fig. **S7**) (Lee *et al*., 2012; Nicaise *et al*., 2013; Wang *et al*., 2020). Our findings therefore illustrate that *Nb*GRP7 is a shared component of PTI and ETI. A parallel can also be drawn with GLK1, which also interacts with the CC domain and potentiates both Rx1-extreme resistance and basal resistance against PVX (Townsend *et al*., 2018). In hindsight, this showed that a single NB-LRR protein can tap into hubs of defence signalling, which fit within a general picture of convergent cell surface-localised and intracellular immune signalling pathways in plant defence.

We demonstrated that *Nb*GRP7 is a pro-immune component of both GpRBP-1 triggered cell death by Gpa2 and extreme resistance by Rx1 in *N. benthamiana* (**Fig. 2a,b,c,d**). *Nb*GRP7 thus adds to the pool of shared co-factors of Rx1 and Gpa2 immunity aside from RanGAP2 (Sacco *et al*., 2007; Tameling & Baulcombe, 2007). These findings suggest that Rx1 and Gpa2 may converge in their use of co-factors and signalling requirements despite their different recognition specificities. This is consistent with sequence exchange experiments showing that the CC-NB of Gpa2 can replace the CC-NB of Rx1 and vice versa while remaining immune receptor function (Slootweg et al 2017). It is striking to note that *Nb*GRP7 overexpression did not affect the autoactive response of Gpa2/Rx1 D460V constructs (Fig. **2a, b**). The D460V mutant is impaired in its MHD motif critical for ADP binding, thereby hampering nucleotide exchange (Moffett *et al*., 2002). We predict that this structural relaxation may override the effect of *Nb*GRP7 needed to surpass the activation threshold. Alternatively, the autoactive response may rely on other host components, which may be rate-limiting for the process but are not regulated by *Nb*GRP7. This in turn reflects a degree of specificity for the role of *Nb*GRP7 in NB-LRR signalling that is reliant on effector-induced changes. However, the precise nature of these changes warrants further investigation.

Despite several optimization attempts, we were unable to demonstrate an effect of *Nb*GRP7 on PVX-CP-triggered cell death (Fig. **2b**). This is fascinating considering that the Rx1-CC and Gpa2-CC domains are highly homologous. As discussed previously, we cannot exclude that GpRBP-1 induced changes can lead to differences between the effect of *Nb*GRP7 on Gpa2 and Rx1 cell deaths. A likely possibility, however, is that the cell death response by Rx1 is too robust. Thus, residual *Nb*GRP7 from overexpression cannot further boost this response. Furthermore, there is accumulating evidence that cell death is dispensable and can be genetically uncoupled from resistance (extensively reviewed in (Künstler *et al*., 2016)). Likewise, extreme resistance by Rx1 to PVX was postulated to be epistatic to cell death (Bendahmane *et al*., 1999). This is further reinforced by structure-function studies of the Rx1-CC, indicating that different surface regions of the domain can be linked to cell death and extreme resistance (Slootweg *et al*., 2018). Thus, we cannot exclude that *Nb*GRP7 may function in regulating extreme resistance while having a limited role in the cell death pathway. Similar outcomes were noted for GLK1, whose overexpression only impacts extreme resistance as well (Townsend *et al*., 2018).

Co-ordinated control of plant NB-LRRs transcripts is key for appropriate defence activation. This has led to an extensive evolution of various molecular checkpoints to fine-tune the dosage of NB-LRRs in the cell. As a corollary, there is ample evidence for splicing, lifetime, and export of mRNAs as a differential response to biotic stress (extensively reviewed in (Lai & Eulgem, 2018)). *At*GRP7 was shown to bind directly to transcripts encoding Pattern-Recognition Receptors *in vivo*, although the consequence of such bindings remains unclear. Here, we demonstrate that overexpressing *Nb*GRP7 directly enhances the transcript and protein levels of intracellular Rx1 (Fig. **4**). Earlier studies performed in potato protoplasts have shown that the extreme resistance response of Rx1 to PVX does not require *de novo* synthesis of defence transcripts (Gilbert *et al*., 1998). In this model, it is, therefore, imperative that a sufficient pool of pre-existing components is available for defence. This puts post-transcriptional regulation at the forefront for regulating Rx1 function. Furthermore, this is in accordance with reports demonstrating that Rx1 transcripts are subject to regulation by 22-nt microRNAs (Li *et al*., 2012). Previous works in potato have shown that modulating Rx1 and Gpa2 transcript/protein abundance directly impacts defence output (Slootweg *et al*., 2017), indicating that exerting control at a post-transcriptional level is important in fine-tuning immunity. We believe that the biological role of plant GRP7s as RNA chaperones fit within this framework. Consistent with this, we observed that the interaction of *Nb*GRP7 and Rx1 localize to speckle-like structures in the nucleoplasm (Fig. **1b)** that are linked to active sites of (post)-transcriptional processing (Spector & Lamond, 2011).

Although the mechanistic basis of how *Nb*GRP7 contributes to Rx1 and Gpa2 at a post-transcriptional level is not fully clear, functional studies with the *Nb*GRP7 R49K/R49Q mutant variants indicate that its RNA binding capacity is involved (Fig.**3** and Fig.**4d, f**). Thus, it will be of interest to determine whether *Nb*GRP7 directly impacts the turnover of Rx1/Gpa2 transcripts as described for *At*GRP7 and FLS2 (Nicaise *et al*., 2013). Imaginably, *Nb*GRP7 could also concurrently regulate multiple targets, for example, defence-transcripts downstream of Rx1. This is reminiscent with the regulation of PR-1 by *At*GRP7 does not involve direct binding to the PR-1 transcript (Hackmann *et al*., 2014). Preliminary data shows that *Nb*GRP7 overexpression upregulates a number of defence marker genes (**Supporting Information** Fig. **S8**). Hereby, it is important to note that our expression analysis did not indicate any nonspecific impacts on the housekeeping gene actin, thus the effect is specific in response to immunity. Future studies should, therefore, aim at elucidating the nature of the immediate cargo bound to *Nb*GRP7.

Altogether, we envision that *Nb*GRP7 belongs to a complex that regulates the transcript homeostasis of the NB-LRR Rx1 and Gpa2, and associated defence genes for immunity (Fig. **5**). By docking to the Rx1/Gpa1-CCs, *Nb*GRP7 arrives in close proximity to other bound interactors in the receptor complex. In the case of Rx1, this may refer to cytoplasmic RanGAP2, which coincides with our observation that *Nb*GRP7 does not share an interacting surface with RanGAP2 on the Rx1-CC (**Supporting Information** Fig. **S4 b, c**) (Sacco *et al*., 2007; Tameling & Baulcombe, 2007). Alternatively, *Nb*GRP7 may be brought in close proximity to other nuclear components like GLK1 and DBCP at the DNA to regulate the function of Rx1 in the nucleus (Fenyk *et al*., 2015; Townsend *et al*., 2018; Sukarta *et al*., 2020). For example, when Rx1 induces transcriptional reprogramming via the activity of transcription factors such as GLK1, *Nb*GRP7 can stabilize the resultant transcripts and thereby, safeguards response outputs. It would, therefore, be fascinating to determine how *Nb*GRP7 would co-operate with existing nuclear interactors of Rx1 and contribute to the transcriptional regulation of downstream immune responses.

**Fig. 5.**
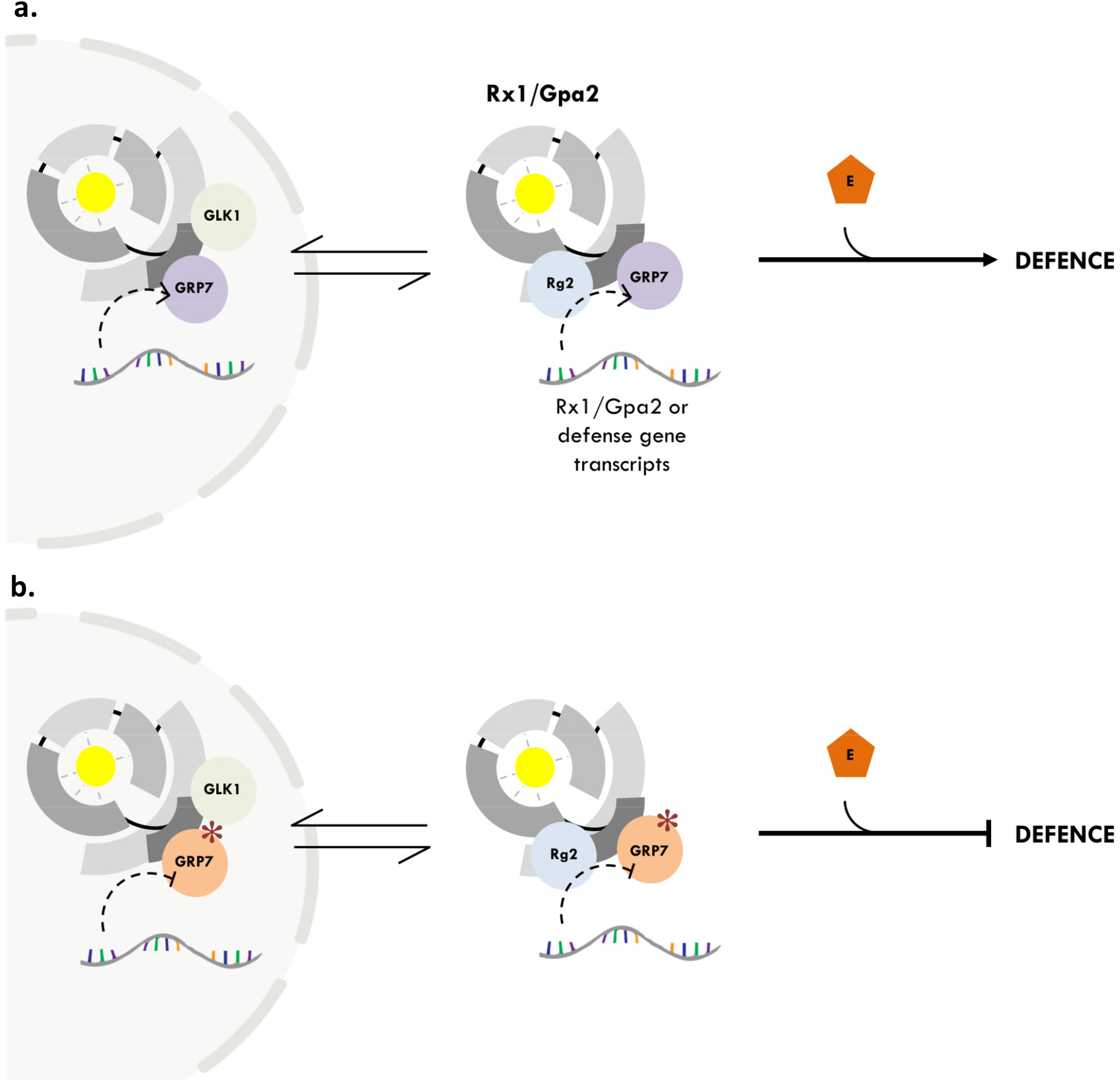
Schematic representation of a working model proposed for the role of *Nb*GRP7 in effector-triggered immunity by Rx1/Gpa2. **a)**. *Nb*GRP7 exists as pre-formed complexes with the receptor proteins in either the nucleus and/or cytoplasm. *Nb*GRP7 is presumed to regulate the transcript and protein levels of Rx1/Gpa2 and/or defence components in the cell through its RNA chaperone activity via a yet undefined mechanism (**curved dashed line**). Presence of the appropriate elicitor (**E**) is recognized in the cytoplasm and triggers a conformational switch in Rx1/Gpa2. Collectively, these changes ensure that a balanced and steady abundance of Rx1/Gpa2 is present to promote an immune response. **b)**. Impairing the RNA Recognition Motif of *Nb*GRP7 (red asterisks) is predicted to compromise its ability to regulate the target transcripts, thereby compromising defences by Rx1/Gpa2.

## ACKNOWLEDGEMENTS

The current work benefits from funding by the Dutch Top Technology Institute Green Genetics (**5CFD051RP**), Dutch Technology Hotel grant and the Dutch Technology Foundation STW and Earth and Life Sciences ALW (STW-GG 14529), which are part of the Netherlands Organization for Scientific Research (NWO). Additionally, we thank Jan-Wilem Borst and Arjen Badder from Wageningen Light and Microspectroscopy Centre for providing imaging facilities and their technical expertise, and Martin Cann and Alexander Llewlyn for their biochemical expertise.

## AUTHOR CONTRIBUTIONS

Conceptualization A.G.; Methodology, O.C.A.S., and Q.Z; Investigation, O.C.A.S., Q.Z., E.J.S., S. B., H.O., R.P., M.M., and V.P.; Writing – Original Draft, O.C.A.S.; Writing – Review & Editing, O.C.A.S., Q.Z., A.G., and G.S; Funding Acquisition, A.G.

## SUPPORTING INFORMATION FIGURE LEGENDS

**Fig. S1**. Summary of data derived from Co-IP/MS analysis of *Nb*GRP7 with Gpa2-CC.

**Fig. S2**. Phylogenetic analysis and multiple sequence alignment of *Nb*GRP7 with other GRP homologs.

**Fig. S3**. BiFC based interaction analysis of *Nb*GRP7 and the CC domain of Rx1 and Gpa2.

**Fig. S4**. Co-IP of *Nb*GRP7 with various subdomains of Rx1 (CC, NB-ARC and LRR) and Rx1 surface-mutant variants.

**Fig. S5** Construct design and silencing efficiency analysis for hairpin silencing of *Nb*GRP7.

**Fig. S7**. The role of *Nb*GRP7 in immunity against PVX-UK3 independent of Rx1.

**Fig. S8**. Ectopic expression of *Nb*GRP7 affects transcript levels of defence marker genes.

**Table S1**. Primers used in the current research as listed according to the assays performed.

**Table S2**. Sequences of *Nb*GRP7 hairpin constructs used in the current research.

