## Supplemental Information for "Glycine-Rich RNA-Binding Protein 7 interacts with and potentiates effector-induced immunity by Gpa2 and Rx1 based on an intact RNA Recognition Motif"

The following Supporting Information is available for this article:

**Fig. S1.** Peptide hits and their locations in the full-length primary sequence of the *Nb*GRP7 homolog identified in the Co-IP/MS screening (shown as bold, underlined sequences). * indicates a ratio of LFQ intensities in the test samples relative to negative GFP control as determined by the label-free MaxQuant algorithm.

| ***Nb*GRP7 (Gpa2-CC-GFP nuclear extract)** | |
|---|---|
| *Peptide hits* | 2 |
| *LFQ Intensity Ratio* (Log*$_{10}$*)* | 1.7 |
| **Peptide Hits** | |
| CFVGGLAWATTDR | |
| NITVNEAQSR | |

MAAEVEYR**<u>CFVGGLAWATTDR</u>**TLGDAFAHYGEVVDSKIINDRETGRSRGF
GFVTFSDEKAMRDAIEGMNGQNLDGR**<u>NITVNEAQSR</u>**GSGGGGGGFGGGR
RREGGYSGGGGYGGGSGGYGGGRREGGYSGGGGGYGGGYGGGRNRGYG
GGYGGGGGDGGSRYSRGGGASEGSWRN

**Fig. S2. a**). **Unrooted Bayesian tree of selected orthologues of NbGRP7.** Selected sequences are indicated by their abbreviated species (e.g. At for *Arabidopsis thaliana*). Support values are indicated at the tree nodes. **b**). Multiple protein alignment of *Nb*GRP7 with characterized paralogs from *Arabidopsis*, potato and pepper generated using the Geneious software ver. 2020.1. Detailed alignment of specific functional motifs and conserved residues are provided in the next row. Orange highlight in the consensus sequence indicates boundaries of the RNA Recognition Motif. Blue triangle indicate the conserved arginine residue in the RRM.

**a.**

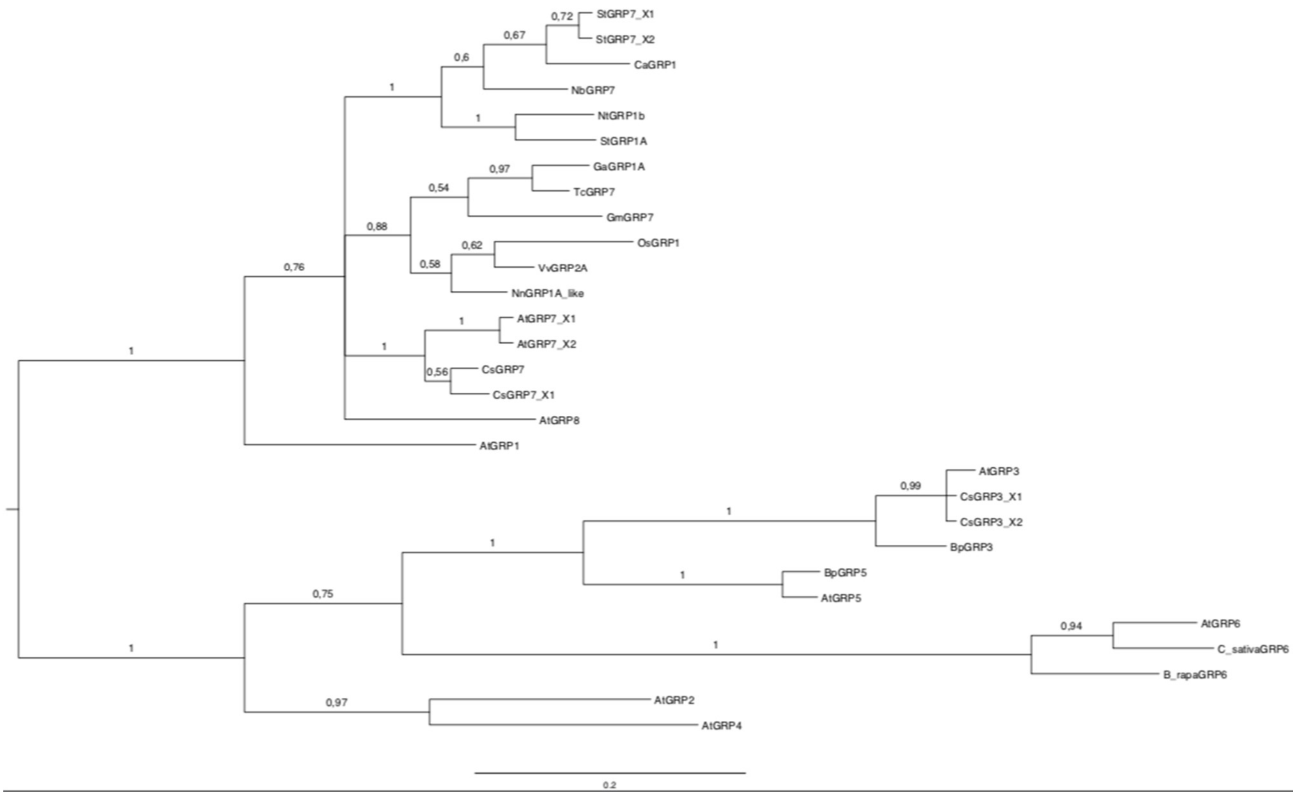

**b.**

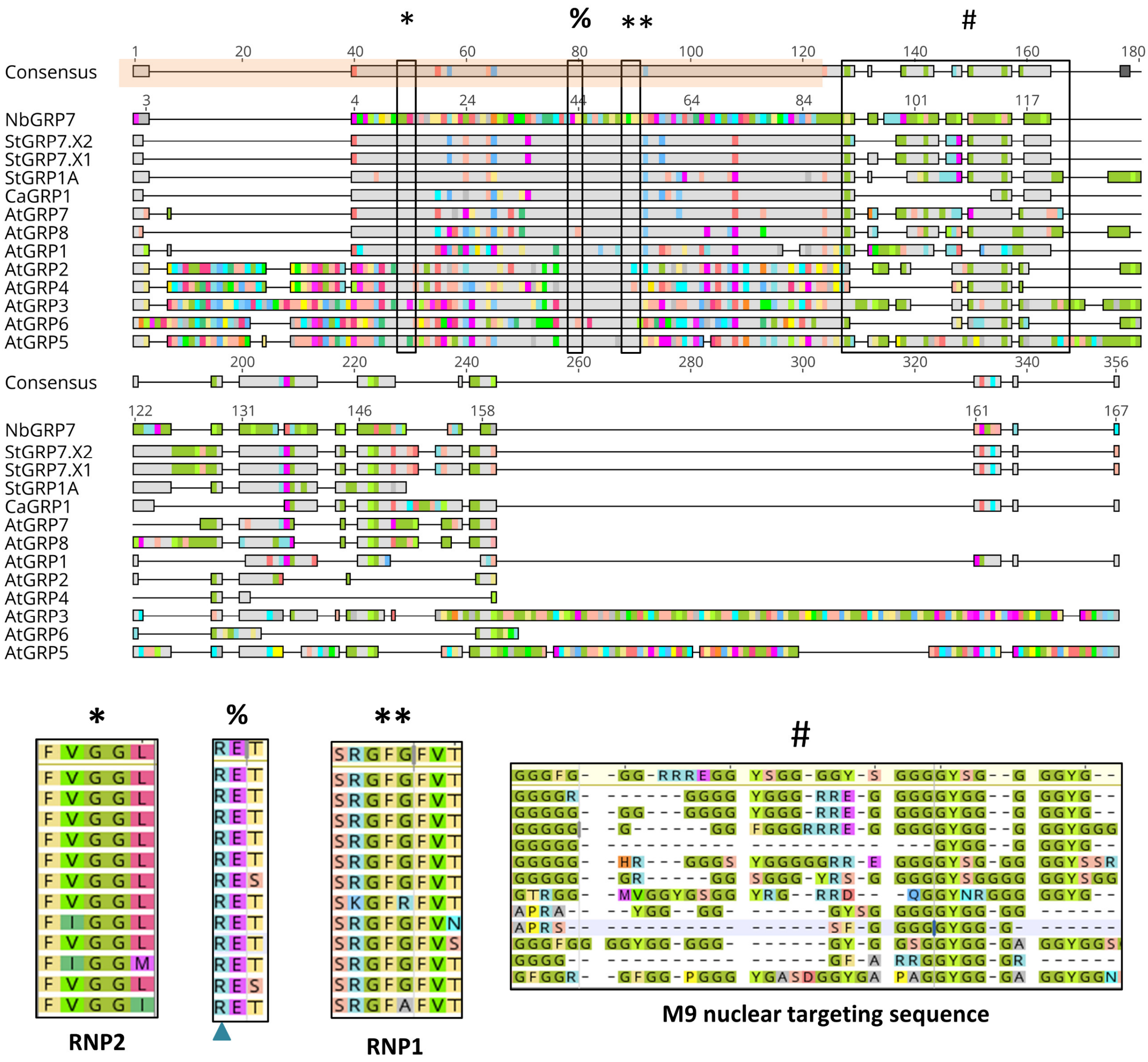
*
%
**
#
Consensus
NbGRP7
StGRP7.X2
StGRP7.X1
StGRP1A
CaGRP1
AtGRP7
AtGRP8
AtGRP1
AtGRP2
AtGRP4
AtGRP3
AtGRP6
AtGRP5
RNP2
RNP1
M9 nuclear targeting sequence

**Fig. S3. BiFC based interaction analysis of NbGRP7 and the CC domain of immune receptors Rx1 and Gpa2. a).** Co-expression of the C-terminal half of the super cyan fluorescent protein SCFP3A fused Rx1-CC or Gpa2-CC (pC:R1 or pC:G1) and the N-terminal half of SCFP3A fused NbGFP7 (pN:NbGRP7) in *N. benthamiana* leaves. Free CFP was used as positive control while co-expression with pC:NSs or pN:GUS were used as negative controls. N, nucleus. C, cytoplasm. NB, nuclear body. Scale bar=10 or 20 µm. Cells were imaged at 2 dpi based on 3 cells. Results are representative of 2 biological repeats. **b). Western blot detection of BiFC constructs.** Co-expression of NbGRP7 and the CC domain of Rx1 or Gpa2. Proteins were extracted at 2 dpi. The N-terminal half of the super cyan fluorescent protein SCFP3A fused proteins (pN:R1, pN:G1, pN:NbGRP7 and pN:GUS) were detected with anti-flag antibody. The C-terminal half of SCFP3A fused proteins (pC:R1, pC:G1, pC:NbGRP7 and pC:NSs) were detected with anti-HA antibody.

**a.**

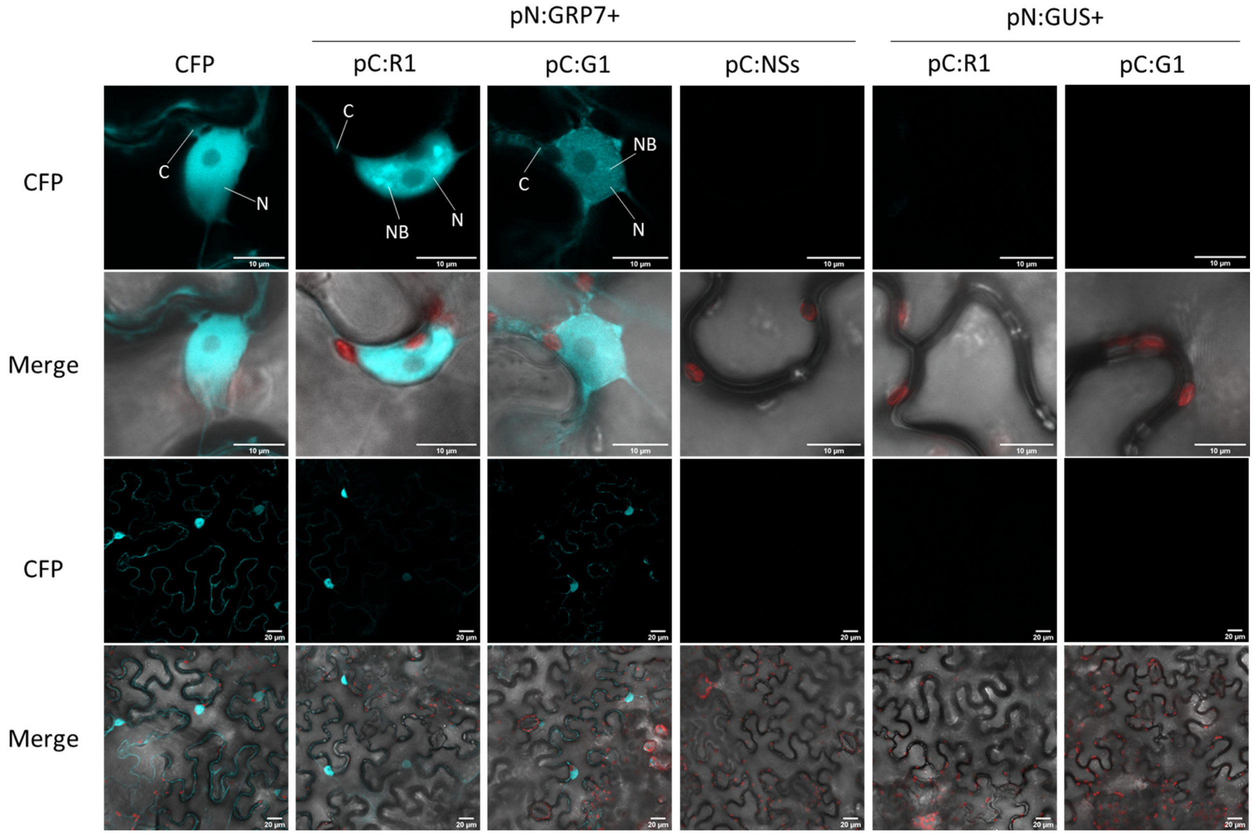

**b.**

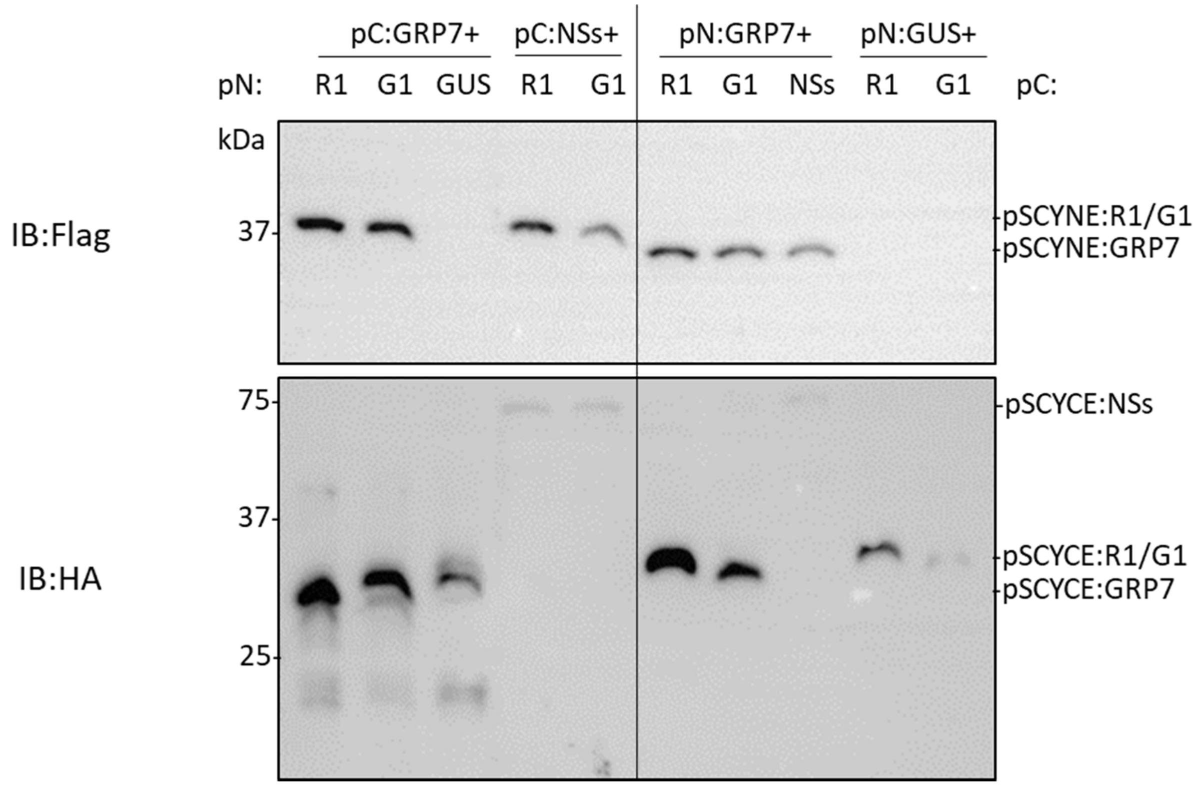
pC:GRP7+
pC:NSs+
pN:GRP7+
pN:GUS+
pN:
R1
G1
GUS
R1
G1
R1
G1
NSs
R1
G1
pC:
kDa
IB:Flag
37
pSCYNE:R1/G1
pSCYNE:GRP7
IB:HA
75
37
25
pSCYCE:NSs
pSCYCE:R1/G1
pSCYCE:GRP7

**Fig. S4. a).** *Nb*GRP7 was used as bait for the Co-IP with the various Rx1 subdomains (LRR, NB-ARC and CC). **b, c)**. Pull-down investigating the potential interaction of *Nb*GRP7 as bait with Rx1 (CC) surface mutants (S1 or S4). In **b**, interaction with the mutated CC domains were investigated whereas in **c**, full-length Rx1 containing these mutations were used. Data shown for each IP is representative of at least two independent repeats. "+" indicates the presence of a construct in the infiltration combination used for immunoprecipitation. In these assays, 4×Myc.GFP of the *Nb*GRP7 fusion construct tends to be cleaved-off under the extraction conditions used. This is indicated by the blue triangle as exemplified in **a**.

**a.**

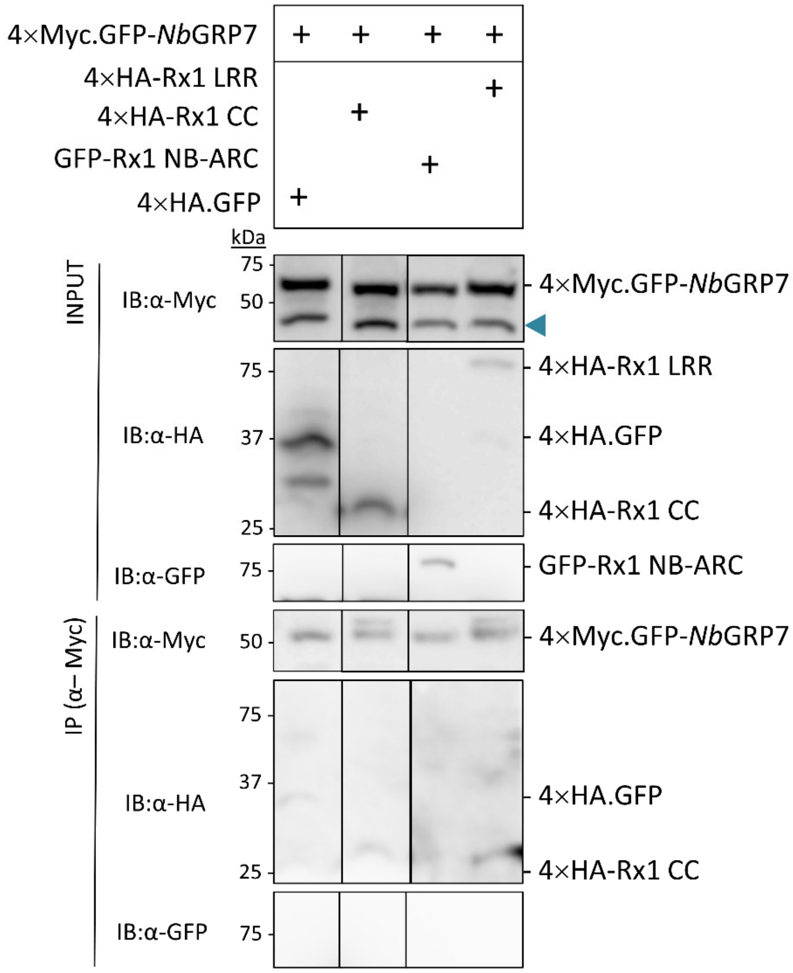

**b.**

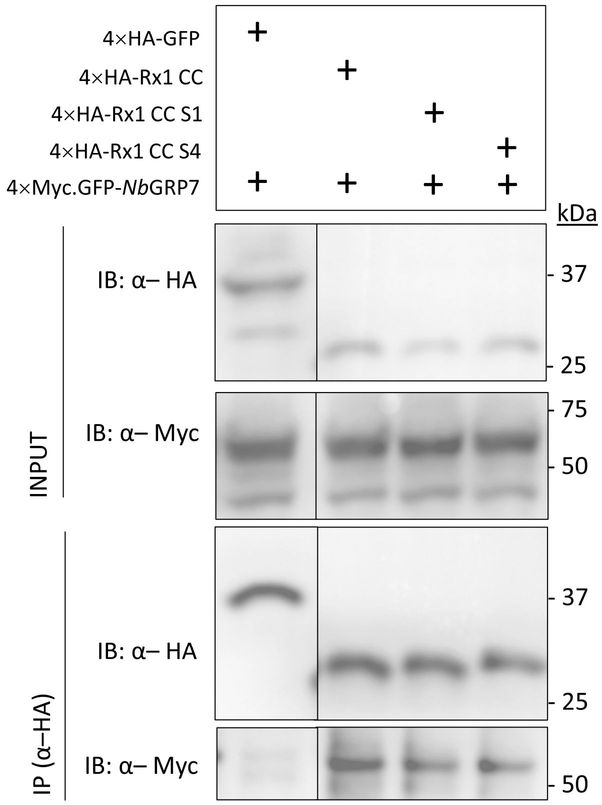
4×HA-GFP
4×HA-Rx1 CC
4×HA-Rx1 CC S1
4×HA-Rx1 CC S4
4×Myc.GFP-NbGRP7
kDa
INPUT
IB: α– HA
37
25
IB: α– Myc
75
50
IP (α–HA)
IB: α– HA
37
25
IB: α– Myc
50

**c.**

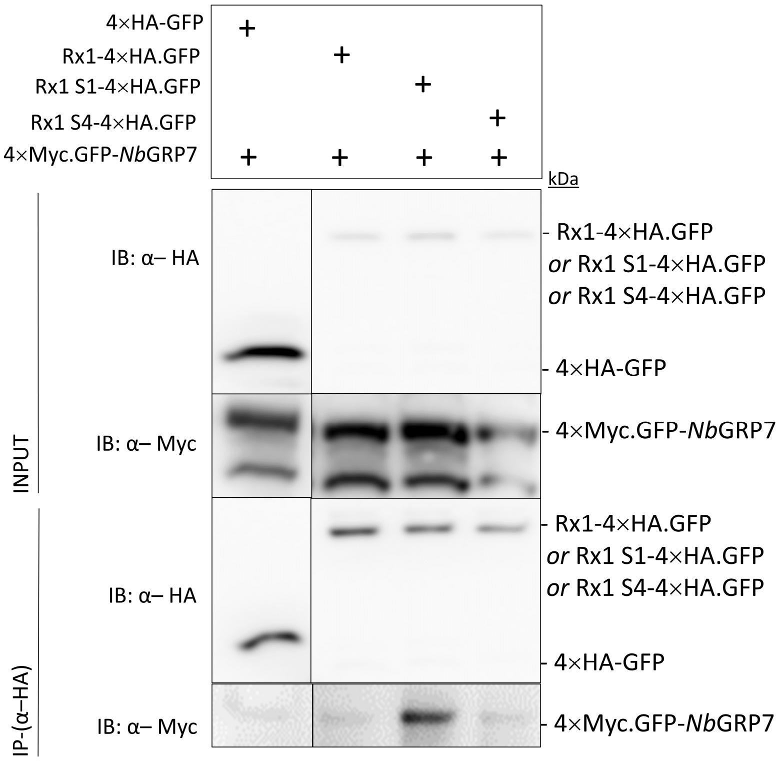
4×HA-GFP
Rx1-4×HA.GFP
Rx1 S1-4×HA.GFP
Rx1 S4-4×HA.GFP
4×Myc.GFP-NbGRP7
kDa
INPUT
IB: α– HA
Rx1-4×HA.GFP or Rx1 S1-4×HA.GFP or Rx1 S4-4×HA.GFP
4×HA-GFP
IB: α– Myc
4×Myc.GFP-NbGRP7
IP-(α–HA)
IB: α– HA
Rx1-4×HA.GFP or Rx1 S1-4×HA.GFP or Rx1 S4-4×HA.GFP
4×HA-GFP
IB: α– Myc
4×Myc.GFP-NbGRP7

**Fig. S5 a)**. Schematic representation of the construct design for hairpin silencing of *Nb*GRP7. Regions corresponding to characteristic domains are indicated above the gene-structure. RRM = RNA Recognition Motif; GRR: Glycine Rich Region. **b)**. Efficiency of silencing on endogenous *Nb*GRP7 transcript is represented as bargraph using RNA extracted from leaf materials harvested at 3 dpi. **c)**. Efficiency of silencing was also tested by immunoblotting experiments of an overexpressed 4×Myc.GFP-*Nb*GRP7 construct in combination with HpGUS control or the two silencing constructs Hp*Nb*GRP7 and Hp*Nb*GRP7_2. CBB-stained membrane of the RuBisCO protein is provided as loading control.

**a.**

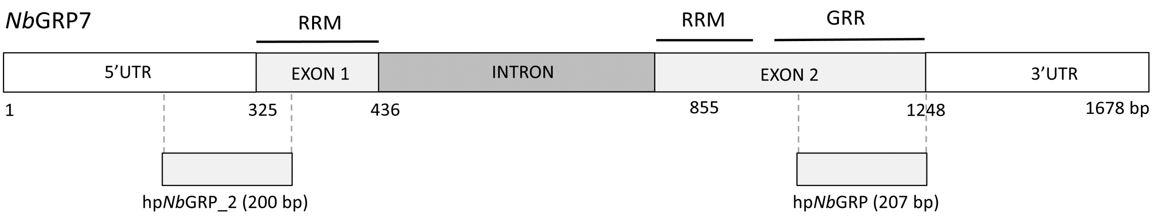

**b.**

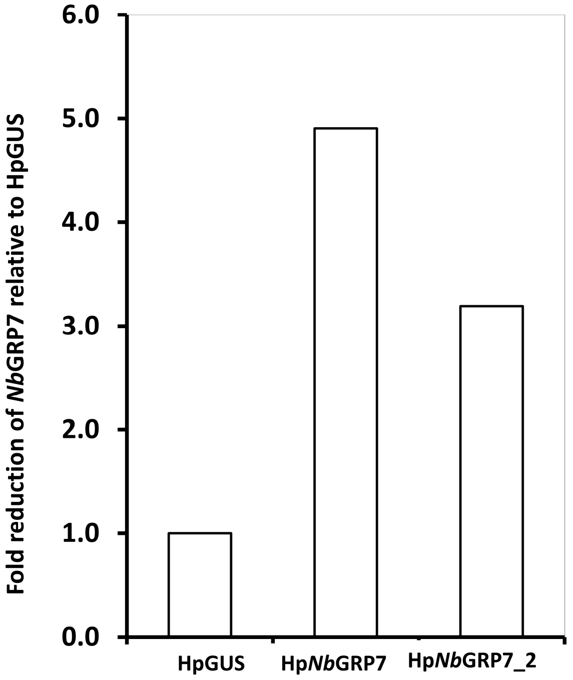

Fold reduction of *Nb*GRP7 relative to HpGUS
6.0
5.0
4.0
3.0
2.0
1.0
0.0
HpGUS
Hp*Nb*GRP7
Hp*Nb*GRP7_2


**c.**

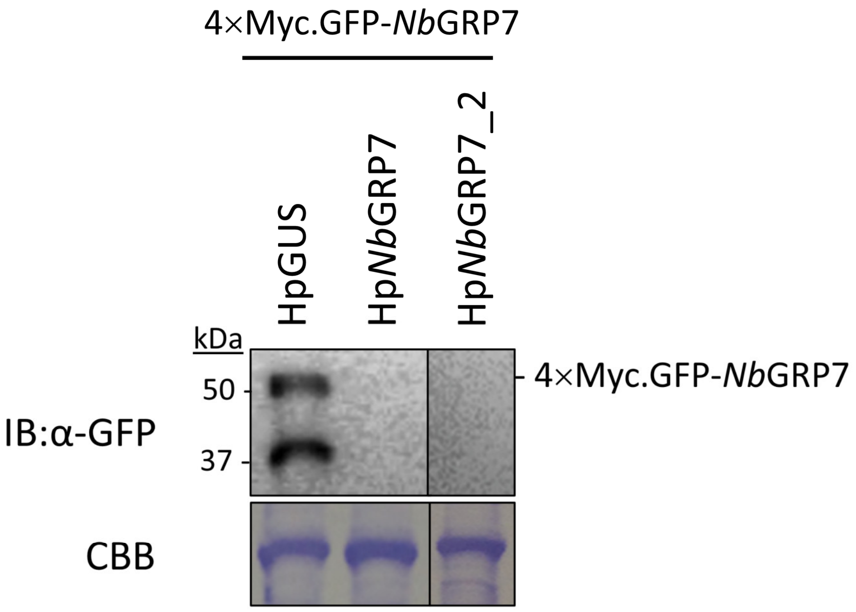

4×Myc.GFP-*Nb*GRP7
HpGUS
Hp*Nb*GRP7
Hp*Nb*GRP7_2
kDa
50
37
IB:α-GFP
CBB
4×Myc.GFP-*Nb*GRP7

**Fig. S6.** Western blot (**a**) and confocal imaging (**b**) of RNA binding mutants of 4×Myc.GFP-*Nb*GRP7 indicating that they are stably expressed *in planta* but localize to different regions in the nucleoplasm. Imaging in **b** was performed at 3 dpi. Scale bar=10 μm.

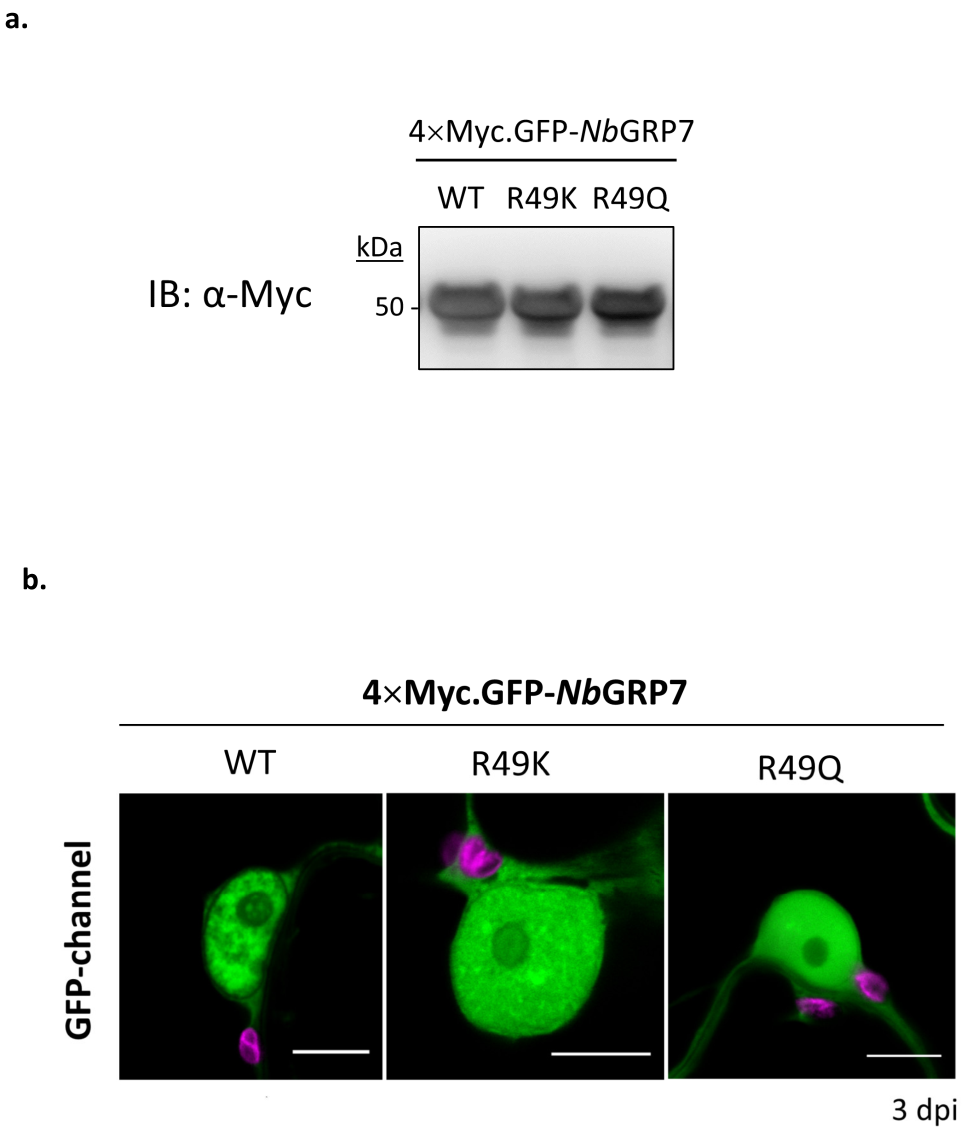

**Fig. S7. *Nb*GRP7 potentiates immunity against PVX-UK3 independent of Rx1.** Boxplots representing absorbance at 405 nm, indicating levels of PVX-UK3 upon transient overexpression of *Nb*GRP7. Bars represent the interquartile range, and the median is indicated by the crossbar. Whiskers show the maximum and minim data points respectively. Data shown is from a single representative experiment (n = 8 samples) with similar results from at least three independent repeats. Significance difference was calculated using Wilcoxon-Signed Rank test with α = 0.05.

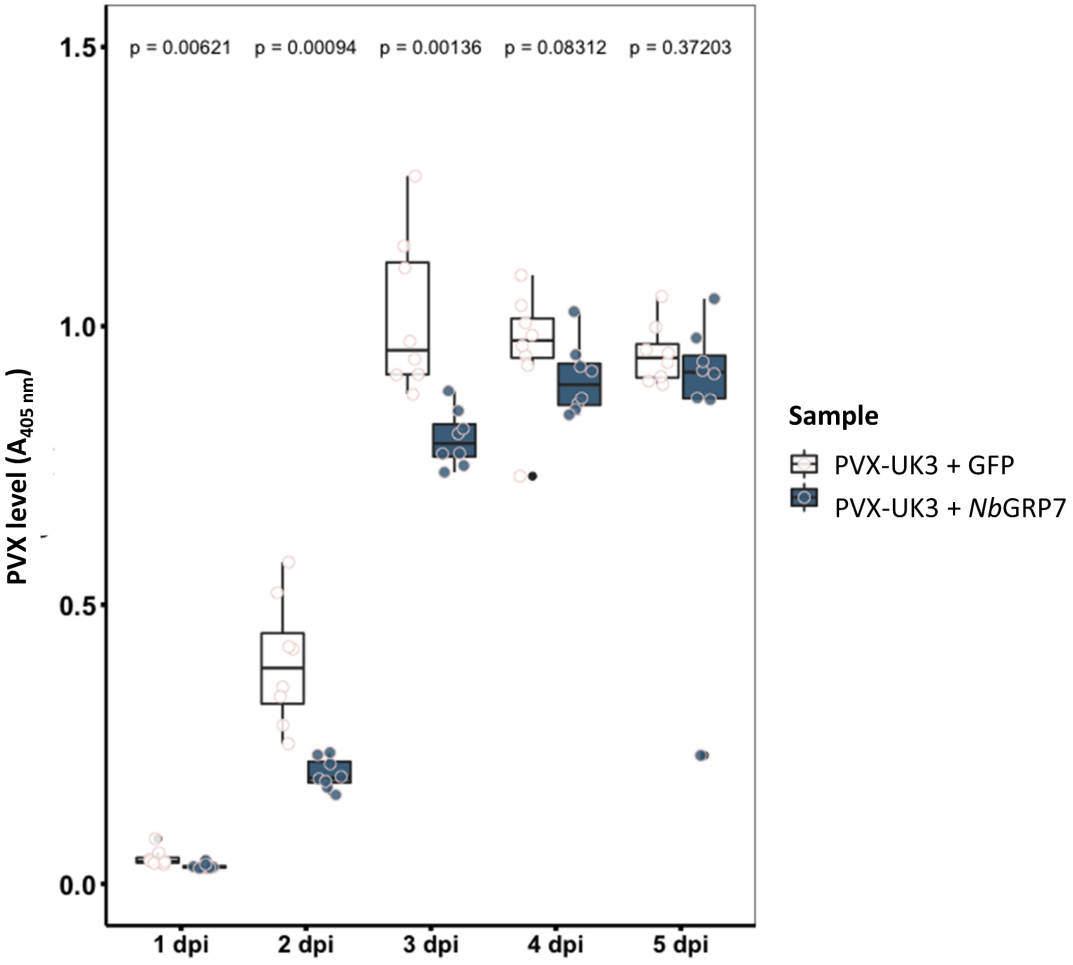

**Fig. S8. Ectopic expression of *Nb*GRP7 affects transcript levels of defence marker genes.** Infiltrated *N.benthamiana* leaves were harvested at 1-5 dpi. Marker genes *HIN1*, *HSR203J* and *bZIP60* were quantified by qPCR. Following normalization to the actin reference gene, relative fold change was determined by comparison of in samples wereby PVX-UK3, Rx1 and *Nb*GRP7 were co-expressed compared to samples of the PVX-UK3 + Rx1 group.

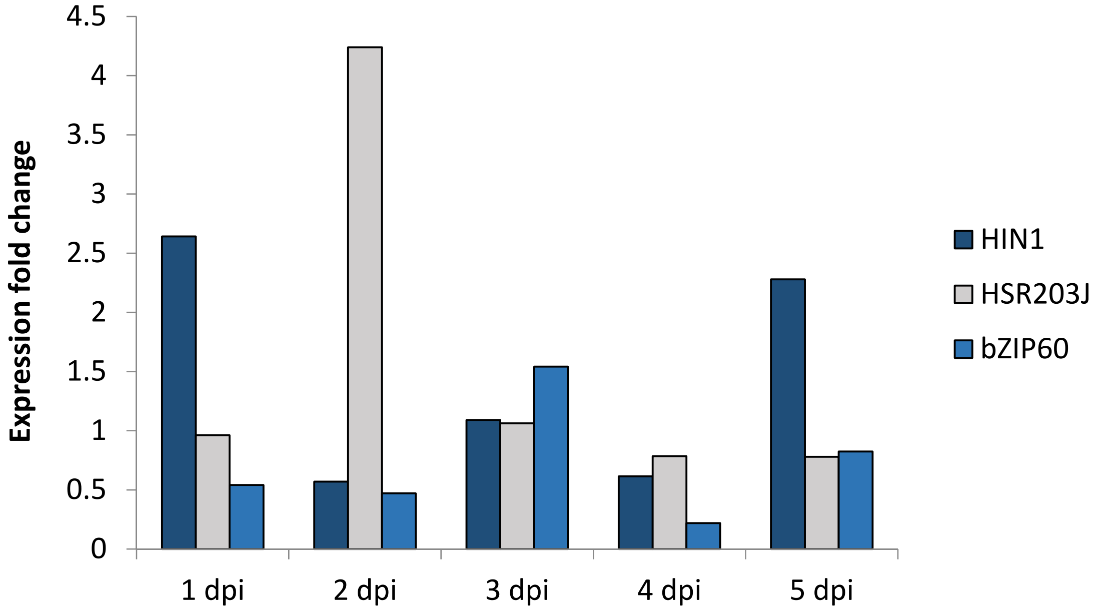

**Table S1.** Primers used in this study as listed according to the assays performed.

| Primer Name | Sequence |
|---|---|
| **Cloning** | |
| *Nb*GRP7 | F: 5'- TGGGCTACCACCGATAGAAC-'3<br>R: 5'- TGCGTCCCTCATAGCTTTCT- '3 |
| *Nb*GRP7 R49K | F: 5'- GAGACTGGAAGATCAAAAGGATTTGGCTTTGTT - '3<br>R: 5'- AACAAAGCCAAATCCTTTTGATCTTCCAGTCTC –'3 |
| *Nb*GRP7 R49Q | F: 5'- GAGACTGGAAGATCACAGGGATTTGGCTTTGTT - '3<br>R: 5'- AACAAAGCCAAATCCCTGTGATCTTCCAGTCTC –'3 |
| **BiFc imaging studies** | |
| Gpa2 CC | F: 5'- CACCATGGCTTATGCTGCTGTTAC - '3<br>R: 5'- CTATAT ATTCTCGGGCTGCTCAAC - '3 |
| Rx1 CC | F 5'- CACCATGGCTTATGCCGCTGTTAC - '3<br>R 5'- CTACATGATATTCTCGGGCTGCTC - '3 |
| *Nb*GRP7 | F 5'- CACCATGGCAGCTGAGGTTGAGTA - '3<br>R 5'- CTAATTCCTCCAGCTTCCCTCGGA - '3 |
| **qPCR analysis** | |
| *Nb*GRP7 qPCR | F: 5'- GAGGATACAGTGGTGGTGGA - '3<br>R: 5'- CTCTTGAGTAGCGGGAACCA –'3 |
| Rx1 qPCR | F: 5'- TTGAGGGAAGCTCGAAACAT - '3<br>R: 5'- ACGACACCAAGCCAATTCTT –'3 |

| | |
|---|---|
| Actin qPCR | F: 5’- CCGAGCGGGAAATTGTTAGG - ‘3 |
| | R: 5’- CACGGATGAGCTGGTCTTTG –‘3 |
| H1N1 qPCR | F: 5’- TTCCGCCACCAGCAAAATC - ‘3 |
| | R: 5’- TTAGGACGAAGAACGAGCCATA –‘3 |
| HSR203J qPCR | F: 5’- AGGCGGCGGCTTTTGTGTCA - ‘3 |
| | R: 5’- GAGAGGTCCCGGAGCCAGAGG –‘3 |
| bZIP60 qPCR | F: 5’- CCTGCTTTGGTTCATGGGCATCAT- ‘3 |
| | R: 5’- AGAAGACCGTGGTTTCTGCTTCGT –‘3 |

**Table S2.** Sequences of hairpin constructs used in this study. Spacer sequence used in the construct for the inverted repeat is highlight in blue. Sites for the restriction enzymes XbaI and BamHI used in sub-cloning are underlined.

| | | |
|---|---|---|
| Hp*Nb*GRP7 | 207 bp | TCTAGAAGGCGGATACAGTGGTGGTGGAGGATACAG<br>TGGTGGTGGAGGATACAGTGGCGGCGGCTATGG<br>AGGTGGAAGACGTGAGGGTGGCTACGGTGGTGGTTA<br>TGGAGGTGGCCGTGACCGTGGATATGGTGGCGGTTA<br>TGGCGGTGGTGATGGTGGTTCCCGCTACTCAAGAGG<br>TGGTGGTGCATCCGAGGGAAGCTGGAGGAATTAACA<br>CTGCACGGTATGCTCCTCTTCTTGTTCATGGTCATGA<br>TCCTTATATGAGCAGGGAAAGTCCAGTTTAGACTTGT<br>AGTTAGTTACTCTTCGTTATAGGATTTGGATTTCTTG<br>CGTGTTTATGGTTTTAGTTTCCCTCCTTTGATGAATA<br>AAATTGAATCTTGTATGAGTTTCATATCCATGTTGTG<br>AATCTTTTTGCAGACGCAGCTAGTAATTCCTCCAGCT<br>TCCCTCGGATGCACCACCACCTCTTGAGTAGCGGGA<br>ACCACCATCACCACCGCCATAACCGCCACCATATCC<br>ACGGTCACGGCCACCTCCATAACCACCACCGTAGCC<br>ACCCTCACGTCTTCCACCTCCATAGCCGCCGCCGCCA<br>CTGTATCCTCCACCACCACTGTATCCTCCACCACCAC<br>TGTATCCGCCTGGATCC |
| Hp*Nb*GRP7<br>_2 | 200 bp | TCTAGAATTCACATCAGCTCTTCTCTTAATTAATTAA<br>CTCTCTCTTGTTTGTTTACATTTATAATCTCTTTCAAG<br>AAAAAAGAGAAAGAAAAACAATGGCAGCTGAGGTT<br>GAGTACAGGTGCTTCGTAGGTGGGCTGGCATGGGCT<br>ACCACTGATAGAACGTTAGGAGATGCTTTTGCTCACT<br>ACGGCGAAGTTGTCGACTCGAAGACACTGCACGGTA<br>TGCTCCTCTTCTTGTTCATGGTCATGATCCTTATATGA<br>GCAGGGAAAGTCCAGTTTAGACTTGTAGTTAGTTAC<br>TCTTCGTTATAGGATTTGGATTTCTTGCGTGTTTATG<br>GTTTTAGTTTCCCTCCTTTGATGAATAAAATTGAATC<br>TTGTATGAGTTTCATATCCATGTTGTGAATCTTTTTGC |

| | | |
|---|---|---|
| | | AGACGCAGCTAGCTTCGAGTCGACAACTTCGCCGTAGTGAGCAAAAGCATCTCCTAACGTTCTATCAGTGGTAGCCCATGCCAGCCCACCTACGAAGCACCTGTACTCAACCTCAGCTGCCATTGTTTTTCTTTCTCTTTTTCTTGAAAGAGATTATAAATGTAAACAAACAAGAGAGAGTTAATTAATTAAGAGAAGAGCTGATGTGAATGGATCC |
| HpGUS | 145 bp | TCTAGACCGCGTCTTTGATCGCGTCAGCGCCGTCGTCGGTGAACAGGTATGGAATTTCGCCGATTTTGCGACCTCGCAAGGCATATTGCGCGTTGGCGGTAACAAGAAAGGGATCTTCACTCGCGACCGCAAACCGAAGTCGGCGGCTTTTACACTGCACGGTATGCTCCTCTTCTTGTTCATGGTCATGATCCTTATATGAGCAGGGAAAGTCCAGTTTAGACTTGTAGTTAGTTACTCTTCGTTATAGGATTTGGATTTCTTGCGTGTTTATGGTTTTAGTTTCCCTCCTTTGATGAATAAAATTGAATCTTGTATGAGTTTCATATCCATGTTGTGAATCTTTTTGCAGACGCAGCTAGAAAAGCCGCCGACTTCGGTTTGCGGTCGCGAGTGAAGATCCCTTTCTTGTTACCGCCAACGCGCAATATGCCTTGCGAGGTCGCAAAATCGGCGAAATTCCATACCTGTTCACCGACGACGGCGCTGACGCGATCAAAGACGCGGGGATCC |
